# Prenatal Stress Dysregulates Resting-State Functional Connectivity and Sensory Motifs

**DOI:** 10.1101/2020.08.26.268359

**Authors:** Zahra Rezaei, Zahra Jafari, Navvab Afrashteh, Reza Torabi, Surjeet Singh, Bryan E. Kolb, Jörn Davidsen, Majid H. Mohajerani

## Abstract

Prenatal stress (PS) can impact fetal brain structure and function and contribute to higher vulnerability to neurodevelopmental and neuropsychiatric disorders. To understand how PS alters evoked and spontaneous neocortical activity and intrinsic brain functional connectivity, mesoscale voltage imaging was performed in adult C57BL/6NJ mice that had been exposed to auditory stress on gestational days 12-16, the age at which neocortex is developing. PS mice had a four-fold higher basal corticosterone level and reduced amplitude of cortical sensory-evoked responses to visual, auditory, whisker, forelimb, and hindlimb stimuli. Relative to control animals, PS also led to a general reduction of resting-state functional connectivity, as well as reduced inter-modular connectivity, enhanced intra-modular connectivity, and altered frequency of auditory and forelimb spontaneous sensory motifs. These resting-state changes resulted in a cortical connectivity pattern featuring disjoint but tight modules and a decline in network efficiency. The findings demonstrate that cortical connectivity is sensitive to PS and exposed offspring may be at risk for adult stress-related neuropsychiatric disorders.

## 1. Introduction

Prenatal stress (PS) is suggested to have had negative effects on 10–35% of children worldwide (Maselko et al., 2015) and is consequently a global public health concern (Kinney et al., 2008; Rubin, 2016). Prenatal stress events include maternal stress and changes in the intrauterine environment during the prenatal period that in turn influence neuroendocrine and immune systems. If PS impacts the maternal neuroendocrine system, it can lead to re-programming of the fetal hypothalamic-pituitary-adrenal (HPA)-axis. For instance, PS down-regulates placental 11β-hydroxysteroid dehydrogenase type 2 (11β-HSD2), an enzyme that metabolizes cortisol (i.e., corticosterone in nonhuman animals) (Jensen Pena et al., 2012; O’Donnell et al., 2012). It may delay GABAergic interneuron development and behavioral inhibition (Lussier and Stevens, 2016) and elevate levels of immune response genes, such as pro-inflammatory cytokines IL-6 and IL-1β (Bronson and Bale, 2014). Depending upon development timing, such alterations can lead to changes in corticogenesis, the birth, migration, and maturation of the neocortical mantle (Bock et al., 2014; Jafari et al., 2020a), with resulting lifelong alterations in brain function (Jafari et al., 2020b) and behavior (Scheinost et al., 2017).

There are a number of potential measures of stress impacts on the brain including changes in sensory-evoked responses (SERs) (Han et al., 2019; McGirr et al., 2020) and alterations in spontaneous resting “default” activity (McCormick, 1999; Raichle, 2010; Ringach, 2009). Resting-state activity is characterized by low-frequency, spontaneous neural activity, and evidence of related fluctuations between some brain areas which shows they are functionally connected (Friston, 2011). In functional connectivity studies, the connectome (or network) consists of regions (or nodes) and their connections (or edges) and provides insights into global brain organization and function in health and disease (Nasiriavanaki et al., 2014). During rest or under anesthesia, functional connectivity has been shown to resemble anatomical (or structural) connectivity (Barttfeld et al., 2015; Grandjean et al., 2017). Resting-state functional connectivity (RSFC) has been used to study changes in brain functional connectivity associated with human psychiatric and neurological disorders (Grandjean et al., 2020; Nasiriavanaki et al., 2014), and mouse analogues of those disorders. RSFC can be assessed using functional neuroimaging techniques in humans and wide-field optical imaging methods in rodents. Among wide-field optical imaging techniques, voltage-sensitive dye (VSD) imaging can provide maps of neuronal activity with mesoscale temporal and spatial resolution and is a convenient way to image spontaneous neural activity (Bermudez-Contreras et al., 2018) and RSFC in mouse models of human neurodevelopmental disorders (McGirr et al., 2020). The brain neocortical connectome is formed during prenatal neurogenesis and postnatal maturation as the brain develops (Scheinost et al., 2017). Consequently, PS as a result of various genetic-, health-, and environment-related factors can influence the development of the connectome (Scheinost et al., 2017; van den Bergh et al., 2018).

Both human and nonhuman studies using neuroimaging techniques have shown that maternal distress during pregnancy is associated with structural and functional alterations in some brain regions, which are reflected in the resting/default mode networks (DMNs) and the attentional networks (van den Bergh et al., 2018). Because of its high temporal resolution, VSD imaging and graph-theoretical representations of imaging results should provide an ideal assessment of the potential effects of PS on the mouse connectome. This was the purpose of the present study. To this end, the neocortical function of adult C57BL/6NJ mice subjected to PS using a repetitive auditory stimulus was investigated using high temporal resolution VSD imaging, and the SERs and the RSFC measures were compared in sensory, motor, and associative cortical regions between the PS and control mice.

## 2. Methods and Materials

### 2.1. Animals

Eighteen female C57BL/6NJ mice between 8-10 weeks of age were individually mated with eighteen male C57BL/6NJ mice in standard pie-shape cages. The animals were given access to food and water ad libitum and maintained on a 12:12-h light:dark cycle in a temperature-controlled breeding room (21 °C) with less than 58±2 dBc room noise level. For the recording of gestational length, a former protocol was followed (Jafari et al., 2017a; Jafari et al., 2018b). When the pups were born, the dams were kept individually with the litters. At the age of 21-23 days, the pups were weaned from their mothers, and one male pup was randomly selected from each litter. Male pups were selected to minimize variation in sex-related susceptibility to the anesthesia used in the acute recording experiments. All experiments and surgeries were performed during the light phase of the cycle under the approved protocols by the University of Lethbridge Animal Care Committee under the regulations of the Canadian Council of Animal Care.

### 2.2. Experimental design

Pregnant C57BL/6NJ mice were randomly assigned into two groups consisting of one PS group and one control group. PS animals were exposed to noise exposure on gestational days (GDs) 12-16 because the corticogenesis process occurs within embryonic days 10-17 in mice, and layers II/III, IV, and V mainly develop during GDs 12-16 (Kolb et al., 2013; Kolb et al., 2012). This timeframe is also in line with the second trimester of human pregnancy, which is a time when substantial human neocortical development occurs (Clancy et al., 2007).

#### 2.2.1. PS procedure

The PS paradigm consisted of an intermittent 3000Hz frequency sound of 90 dB for 1s duration and 15s inter-stimulus interval and lasted for 24hrs starting at 8:00am (Jafari et al., 2017b; Jafari et al., 2019). This tonal frequency was applied because it is audible to mice (Heffner and Heffner, 2007), and is relatively similar to environmental noises, which are largely made up of low to mid-frequency tones (Chang et al., 2014). The intermittent presentation of sound stimuli, as well as 24hrs rest after every exposure also were used to prevent stimulus-induced hearing loss (White et al., 1998). On GDs 12, 14, and 16, pregnant mice (n=6) in groups of two to three in their standard cage were moved to a sound chamber specified for the PS group. A speaker, which emitted the noise stimulus was placed inside the cage. The sound pressure level was monitored daily inside the cage without an animal (Tektronix RM3000, Digital Phosphor Oscilloscope). A similar procedure was applied for the control pregnant mice (n=12). A silent speaker, however, was placed inside the cage, and no noise exposure was given.

#### 2.2.2. Plasma corticosterone assay

A day before each animal surgery, blood was collected from the submandibular vein, between 7:30 to 8:30am. Approximately 0.1ml of blood was taken in heparin-coated tubes. The tubes were centrifuged at 6000rpm at 4°C for 15min to gather the plasma. Collected plasma samples were stored at −80°C and then analyzed using previously described methods (Jafari et al., 2018a; Jafari et al., 2018b).

#### 2.2.3. Surgery

For imaging the neocortex, an acute craniotomy was performed for the PS (n=6) and control (n=12) offspring at age 2-3 months. Mice were given an acute 7×6mm unilateral craniotomy and the dura was removed (bregma +2.5 to -4.5mm, lateral to the midline 0 to 6mm for VSD imaging experiments as previously described (Kyweriga et al., 2017)). Isoflurane (1.0–1.5%) was used to anesthetize mice both for induction and during surgery. All imaging sessions were performed on anesthetized mice after their craniotomy surgery. Although anesthesia may potentially compromise brain recording, previous studies (Mohajerani et al., 2013) have reported no significant differences in sensory-evoked VSD response parameters as well as patterns of spontaneous activity in anesthetized vs quiet awake mice. Moreover, the isoflurane inhalation that was used has been shown to leave the stress response (corticosterone level) intact (Wu et al., 2015). Nevertheless, the concentration of isoflurane (0.5-0.7%) was reduced during the period of data collection. Mice were head-fixed during imaging, in which the skull being fastened to a steel plate while they were posed on a metal plate onto the stage of the upright microscope (Mohajerani et al., 2010). A heating pad was used to preserve the body temperature at 37°C during surgery and imaging sessions.

#### 2.2.4. VSD imaging

Wide-field optical imaging of summated cortical voltage activity was used to capture the mesoscale dynamics of the cortex. For in vivo VSD imaging, the dura was carefully removed within the craniotomy window. The dye RH-1691 (Optical Imaging, New York, NY, USA) (Shoham et al., 1999) was dissolved in HEPES-buffered saline (0.5 mg/ml) and applied to the exposed cortex for 30-40min (Mohajerani et al., 2010). The unabsorbed VSD was washed and imaging begun almost 30min later. Because respiration can lead to movement artifact, the brain was covered with 1.5% agarose made in HEPES-buffered saline and sealed with a glass coverslip to minimize that effect. A charge-of front-to-front video lenses (8.6 × 8.6mm field of view, 67μm per pixel). VSD was excited using coupled device camera (1M60 Pantera, Dalsa, Waterloo, ON, Canada) and an EPIX E8 frame grabber with XCAP 3.8 imaging software (EPIX, Inc., Buffalo Grove, IL, USA) was used to capture 12-bit images with 6.67ms temporal resolution. Collected images were taken through a microscope composed a red LED (627nm center, Luxeon Star LEDs Quadica Developments Inc., AB, Canada) and excitation filter (630±15nm, Semrock, NY, USA). VSD fluorescence was filtered using a 673 to 703nm band-pass optical filter (Semrock, NY, USA). The imaging was focused on the cortex to a depth of ∼1mm to avoid distortion of the signal given the movement of superficial blood vessels.

#### 2.2.5. Recording spontaneous and evoked activity

Spontaneous activity was recorded with 6.7ms temporal resolution (150Hz) prior to recording evoked activity, to prevent sensory stimulation from altering the characteristics of spontaneous activity (Han et al., 2008). Each epoch of spontaneous activity recording lasted for 67 sec, and between 6 to 10 epochs were recorded for each animal. Between each epoch, HEPES-buffered saline was added to the agarose to keep the brain moist. After the spontaneous activity was recorded, the evoked activity procedures were undertaken. Cortical evoked responses forelimb, hindlimb, whisker, visual, and auditory stimuli were measured. A thin needle (0.14mm) was inserted in the paws for hindlimb and forelimb stimulation, which delivered a current of 0.2-1mA for 1ms. For whisker stimulation, a single whisker was attached to a piezoelectric device (Q220-A4-203YB, Piezo Systems, Inc., Woburn, MA, USA) and stimulated by a 1ms square pulse. Visual stimulation was provided through a 1ms pulse of combined green and blue light. For auditory stimulation, a 12kHz pure-tone at 80dB was presented using a Tucker-Davis Technologies (TDT) RX6 once the animal was sitting in a sound-proof booth. The speaker (TDT, ES1 electrostatic loudspeaker) was calibrated to emit a uniformly distributed sound stimulus. Sensory stimulations were separated by an interval of 10s. Stimulated trials lasted for 1s, and the stimuli were presented after passing one-third of the time of that trial. For each animal and each stimulus, between 10 to 20 evoked trials were recorded.

### 2.3. Data analysis

#### 2.3.1. VSD processing

Six to twenty stimulation trials were averaged to decrease the effect of spontaneous change of brain states. These trials were used for the normalization of the stimulated data. Using MATLAB® (Mathworks, Natick, MA, USA), VSD responses were calculated as a percentage change relative to baseline (ΔF/F_0_ × 100%). The baseline was estimated for both no-stimulation trials and spontaneous activity, using the “locdetrend” function from the Chronux toolbox (Bokil et al., 2010). Temporal low-pass filters with cutoff frequencies of 30Hz and 6Hz were respectively used for evoked and spontaneous recordings. To get a smoother signal, a spatial Gaussian filter with sigma equal to 1µm also was applied to both evoked and spontaneous recordings.

#### 2.3.2. ROI identification

The neocortex was divided into 19 regions including primary sensory and motor areas (motor barrel cortex (mBC), motor forelimb (mFL), motor hindlimb (mHL), primary hindlimb (HLS1), primary forelimb (FLS1), primary lip (LpS1), primary barrel cortex (BCS1), primary visual (V1), primary auditory (A1)), supplementary areas (secondary barrel (BCS2), secondary hindlimb (HLS2A and HLS2B), secondary forelimb (FLS2), lateral secondary visual (V2L), medial secondary visual (V2M), anterior segment of the secondary motor (aM2), posterior segment of the secondary motor (pM2)), and association areas (parietal association (ptA) and retrosplenial cortex (RSC)). To specify cortical regions, sensory stimulation was used to set coordinates for the primary and secondary somatosensory areas (HLS1, HLS2A, HLS2B, FLS1, FLS2, BCS1, BCS2, V1, and A1). The coordinates of other areas were set relative to these sensory areas’ coordination, using a stereotaxic atlas(Paxinos, 2001). We defined the regions of interest (ROI) as a 5×5-pixel area around the specified coordinates and the time series associated with each ROI was the average of ΔF/F_0_ across all pixels within the ROI.

#### 2.3.3. Sensory evoked responses

Evoked responses were characterized by peak amplitude, the area under the peak, and time to peak (rise time). The peak of the evoked responses was set as the maximum value from the onset of the stimulus to 200ms thereafter, as long as the value was at least two standard deviations above the baseline. The area under the peak also was defined from the onset of the stimulus to 200ms afterward. Finally, the rise time was determined by subtracting the response onset time from the peak time. For all quantifications, the baseline was defined as the mean of the evoked signal from 300ms to 10ms before the stimulus.

#### 2.3.4. Correlation and network analysis

Correlation matrices were created based on the zero-lag correlation between spontaneous VSD ΔF/F_0_ signals of every couple of 19 cortical ROIs. For network analysis, both full and partial correlations were employed to respectively characterize the global and local structure of the networks. The brain connectivity toolbox (Rubinov and Sporns, 2010) was used to calculate network measures including global efficiency (the average of inverse shortest path length), characteristic path (the average shortest path length between all pairs of nodes in the network), clustering coefficient (the extent that a node’s neighbors are neighbors of each other), nodes betweenness centrality (the fraction of all shortest paths in the network that contains a given node), modularity (the extent a network is divided into delineated communities), and participation coefficient (the extent that a node is restricted to its community). Negative weights were considered in the algorithms we used for measuring modularity and participation, meaning that a portion of these measures came from negative connections. Overall, all network measures were individually found for each animal and then averaged across animals for a group-level comparison.

To find out the community (modularity) structure of the networks, the community Louvain algorithm was employed (Blondel et al., 2008). To adjust the size of the communities, a similar value of resolution parameter (0.2) was used for both control and PS networks. After finding the modularity structure for each animal, it was required to obtain consensus communities over all animals in each group to do a group-level comparison. Hence, first, the concept of multilayer networks (Betzel et al., 2019) (i.e., network layers represent individual subjects’ networks) was applied. Then, the agreement matrix (Betzel et al., 2013), whose elements indicate the number of times any two vertices were assigned to the same community, was obtained. Eventually, through successively applying the Louvain algorithm on the agreement matrix (Lancichinetti and Fortunato, 2012), a consensus partition for networks of both groups was created. Nodes whose degree was in the 80th percentile and had a participation coefficient greater than 0.3 were chosen as connector hubs (Sporns et al., 2007).

For thresholding, we avoided weight thresholding, applying a fixed threshold value to all networks, which leads to a different number of edges in the control and PS networks and is characterized as the main source of group differences (van Wijk et al., 2010). The effect of statistical significance threshold, and proportion/density thresholding, which preserves the same number of edges in all networks by keeping only the strongest X percent of connections (Buchanan et al., 2020; Fornito et al., 2016), is explained below.

##### Correlation analysis

The statistical significance threshold did not affect any of the correlation coefficients as all the p-values were much smaller than the significance level, even after correcting for multiple hypotheses testing (all p-values were less than 10^-10^ even for the smallest correlation values, which were of the order of 0.1). As the histogram in Fig. 3B shows, the connectivity strength distributions were significantly different between groups. To test the hypothesis that this difference was the main reason for the group differences, keeping the same value of connections (i.e., preserving the same histogram of connectivity strength), we generated an ensemble of 100 shuffled connectivity matrices. Supplementary Table S5 shows the result of comparisons. We also examined proportion thresholding. Starting from fully connected networks, as we raised the threshold, the network measures became less different between the two groups. When reaching the backbone of the networks (10-15% of all links), none of the network measures appeared significantly different. We would like to stress, however, that the efficiency of the PS network remained significantly lower than the one of the control network at different threshold levels if at least 30% of all links are kept.

##### Partial correlation analysis

The statistical significance threshold was applied to reduce the possibility of having nodes connected by chance. With this threshold the partial correlation networks became slightly sparser (e.g., removal of 3-11% and 6-16% of the links at the significance level of 0.05 and 0.001, respectively). We also investigated the effect of density thresholding on differences in network topologies. Modularity and the number of connector hubs were significantly higher in the PS networks until 50% (86 out of 171) and 20% (34 out of 171) of links were preserved, respectively. Although PS networks consistently had more connector hubs than control networks at different threshold levels, by keeping less than 70% of connections, the regions emerging as hubs became more variable reflecting stronger statistical fluctuations due to the small network size. Therefore, we first applied a significance threshold to partial correlations and then to equalize network sizes, we kept the strongest 80% of connections, which led to 138 out of 171 possible connections in all networks. Due to the subtraction of all possible confounding correlations, the partial correlation coefficient between two regions can take negative values (Supplementary Fig. 4). These negatively weighted edges may be neurobiologically relevant in functional neuroimaging (Kazeminejad and Sotero, 2020; Sporns and Betzel, 2016). Because there is no consensus regarding the treatment of negatively signed edges, negative correlations were not removed in this study for two main reasons: 1) We did not perform global signal removing (common in fMRI studies (Schwarz and McGonigle, 2011)) in the pre-processing step, which could give rise to anti-correlations. Hence, it is more likely that negative correlations are neurobiologically relevant in our study; 2) Negative weights comprised a considerable portion (36-44%) of all connections both in control and PS networks.

#### 2.3.5. Spontaneous sensory motifs

Sensory motifs are periodic waves of spontaneous activity that resemble evoked responses to sensory stimuli. A template matching technique was used to identify these motifs as previously described (Mohajerani et al., 2013). For each sensory modality, the template was identified using the first three frames after the onset of evoked activity and the Pearson correlation coefficient was taken between templates and the spontaneous activity. Then, frames of the spontaneous activity with a greater correlation coefficient than a given threshold were considered as a ‘match’ (i.e., sensory motif). To determine a correlation threshold for the template matching process, first, a range of thresholds was defined by summing the mean correlation between each sensory template and spontaneous activity with a standard deviation (SD) in steps of 0.1 SD. Then, the number of “matches” obtained from each threshold versus the threshold range was plotted (Supplementary Fig. 5). The threshold level was set where the percentage of “matches” reduced to 36% (e^-1^) of its initial value (Mohajerani et al., 2013). This criterion was applied to all sensory templates individually. The overall threshold level, finally, fell around mean correlation+1.6SD for the PS group and mean correlation+1.7SD for the control group. The minimum interval between two consecutive motifs to consider them as separate events was set at 200ms.

To study the temporal order of sensory motifs within the spontaneous activity, all possible non-repetitive combinations of motifs using the combination formula 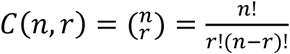, which gives the number of possible r-combinations from a larger sample of n elements, were found. Given five different sensory motifs, 20 double (e.g., auditory-forelimb (AF), …), 60 triple (e.g., auditory-forelimb-visual (AFV), …), 120 quadruple (e.g., auditory-forelimb-visual-whisker (AFVW), …), and 120 quintuple (e.g., auditory-forelimb-visual-whisker-hindlimb (AFVWH), …) non-repetitive combinations were possible. To find these sequences, the frames of spontaneous activity matched with sensory templates were identified. Then, a combination with a certain temporal order of motifs was explored. To name a combination of motifs as a sequence, an upper limit of 1s interval between consecutive motifs was set. To verify that the order matters, the results of the experiment were compared with randomly generated sequences. To make random sequences, the whole string of sensory motifs within a spontaneous recording was temporally shuffled (1000 times), whereas the total number of each sensory motif was kept the same as its actual count.

#### 2.3.6. Statistical tests

We utilized a two-sided (non-directional) Wilcoxon rank-sum test, equivalent to a Mann-Whitney U-test, for cross-sectional analysis. Both full and partial correlation analyses were performed based on the linear dependence of variables using the Pearson correlation coefficient. P-values for correlational analysis were computed using a Student’s t distribution for a transformation of the correlation. This was exact for normal data but was a large-sample approximation otherwise. All statistical analyses were performed using MATLAB at a significance level of 0.05. The Benjamini & Hochberg (BH) procedure was applied for controlling false discovery rate (FDR) (Benjamini and Hochberg, 1995) wherever multiple hypotheses were tested. Error bars and ± ranges represent the standard error of the mean (SEM). Asterisks indicate \**p* < 0.05, \*\**p* < 0.01, \*\*\**p* < 0.001.

##### Data availability

Processed data and codes are available in Mendeley Dataset https://data.mendeley.com/datasets/xfkhsnjcmg/draft?a=462d091e-5458-4b80-9952-a61cc7c08afb.

## 3. Results

### 3.1. Elevated corticosterone levels

The timeline of the experiment is shown in Fig. 1A. The basal corticosterone level was more than four-fold higher in the PS group compared with the control group (p≤ 0.001, Fig. 1B).

**Fig 1.**
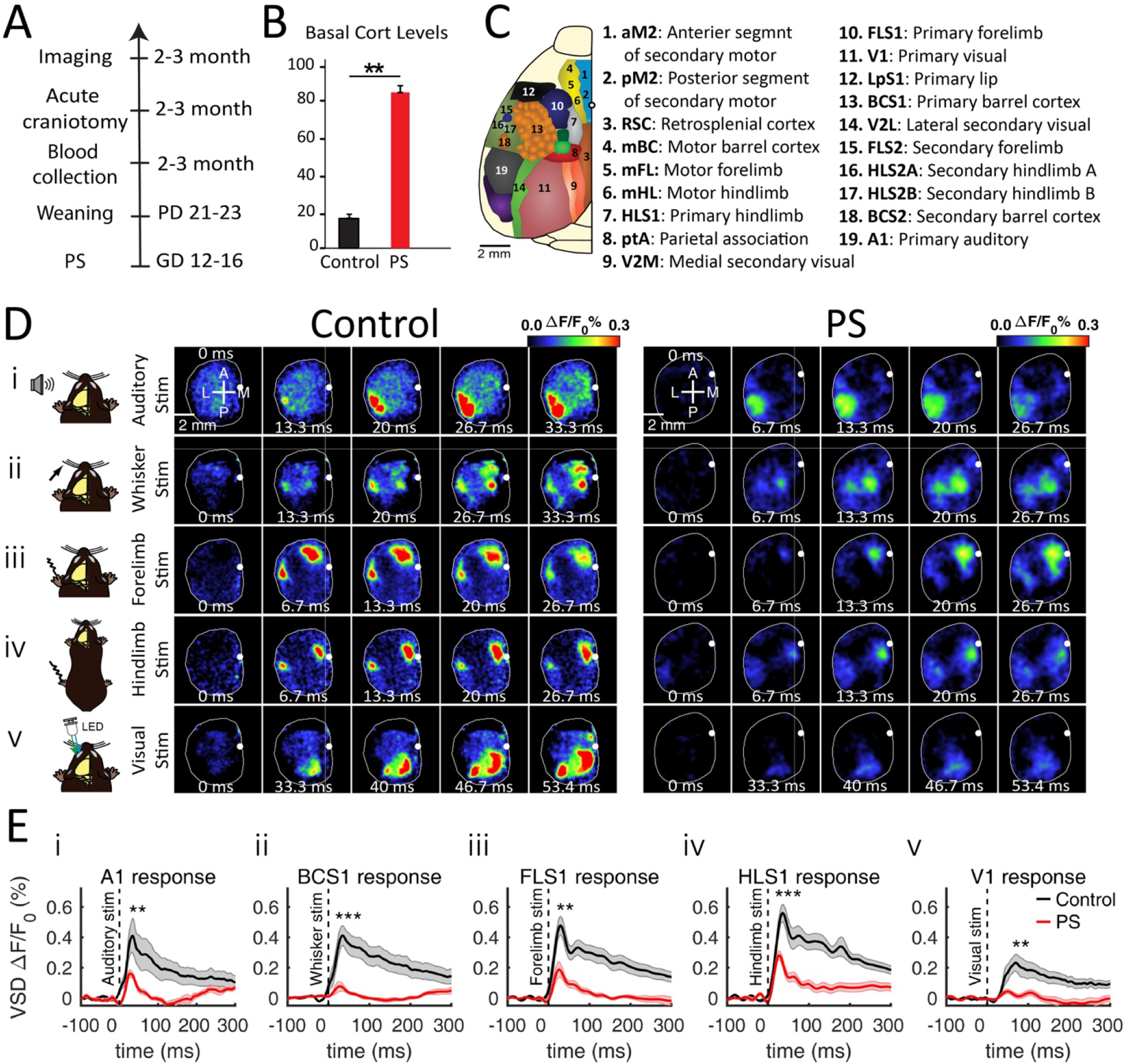
The decline of evoked responses to sensory stimuli in PS mice. **A**) Timeline of the experiment. Pregnant mice were exposed to noise stress on GDs (gestational days) 12-16. Their pups were weaned at the age of 21-23 days (PD, postnatal day). Blood was collected from pups at age 2-3 months (Cort, corticosterone). A day after, an acute craniotomy was performed for both groups. The VSD imaging was conducted a week after. **B**) The basal corticosterone level of PS mice was significantly increased compared to the control mice. **C**) Unilateral craniotomy represents imaged cortical regions. **D**) Photomicrograph of the wide unilateral craniotomy with bregma marked with a white dot in each image. Patterns of cortical activation are shown in a mouse anesthetized with isoflurane (0.5%) after (i) auditory, (ii) whisker, (iii) forelimb, (iv) hindlimb, and (v) visual stimulations with a light-emitting diode (LED) for a control and a PS mouse. These VSD imaging montages show the frame of the stimulus (0 ms) followed by four other frames starting from the onset of the evoked response. The first image from the left in the first row indicates the anterior (A), posterior (P), medial (M), and lateral (L) directions. **E**) Evoked VSD responses to five sensory stimuli. The shaded region in all diagrams shows SEM (standard error of the mean). The asterisks point to a significant difference in the area under the peak. (i) Response of the A1 region after the auditory stimulus (df=13, sum of ranks=96, p=0.003); (ii) response of the HLS1 region after the hindlimb stimulus (df=13, sum of ranks=98, p=8×10^-4^); (iii) response of the FLS1 region after the forelimb stimulus (df=13, sum of ranks=95, p=0.005); (iv) response of the BCS1 region after the whisker stimulus (df=13, sum of ranks=99, p=4×10^-4^); and (v) response of the V1 region after the visual stimulus (df=13, sum of ranks=93.5, p=0.008). The number of mice: n=9 control, n=6 PS. Asterisks indicate *p<0.05, **p<0.01, or ***p<0.001.

### 3.2. Reduced sensory-evoked amplitudes

In the control group, all modes of sensory stimulation (hindlimb, forelimb, whiskers, visual, or auditory systems) resulted in consensus patterns of cortical depolarization (Fig. 1C and 1D) consistent with previous findings (Mohajerani et al., 2013). Sensory stimulation led to the activation of primary sensory areas as well as “islands” of response within functionally related areas such as primary motor regions (mFL, mHL, mBC), secondary sensory regions (FLS2, HLS2, BCS2, V2) or cortical areas along the mid-line (RSC, cingulate, and secondary motor) (Fig. 1D). For all sensory modalities, the activity initially appeared in the primary cortices and then spread to secondary somatosensory cortices. The montage of VSD images in Fig. 1D illustrates a reduction of cortical activation and localization of response to all stimuli in the PS group relative to the control mice.

Fig. 1E shows the evoked response of primary cortical regions to each corresponding stimulus. The PS mice showed a declined VSD response (i.e., ΔF/F_0_) in all cortical regions compared to the control group. To statistically verify this group difference, the peak amplitudes (V1: df=13, sum of ranks=92.5, p=0.012; A1: df=13, sum of ranks=95, p=0.004; BCS1: df=13, sum of ranks=99, p≤0.0001; FLS1: df=13, sum of ranks=94, p=0.007; HLS1: df=13, sum of ranks=95, p=0.004; two-tailed rank-sum test) and the area under the peak (V1: df=13, sum of ranks=93.5, p=0.008; A1: df=13, sum of ranks=97, p=0.001; BCS1: df=13, sum of ranks=99, p≤0.001; FLS1: df=13, sum of ranks=95, p=0.004; HLS1: df=13, sum of ranks=98, p≤0.001; two-tailed rank-sum test) were compared between the two groups. To quantify evoked responses within wider cortical regions, the VSD responses were compared at 19 cortical ROIs corresponding to Fig. 1C (see Methods). The statistically quantified responses of all other cortical regions in terms of peak amplitude, area under the peak, and rise time of the response were shown in Supplementary Materials (Supplementary Fig. 1, 2, and 3; and Tables 1, 2, and 3).

### 3.3. Group difference comparisons using intracortical functional connectivity matrices

Mice in both control and PS groups exhibited dynamic, spatially complex patterns of spontaneous cortical activity. Zero-lag correlation analysis was used to quantify the effect of PS on intracortical functional connectivity. Correlation matrices in Fig. 2A show the average of animals per group. Whereas most of the regions were highly correlated, the overall correlation strength was reduced in the PS compared to the control group (MeanCorr_Control_= 0.8±0.02 vs. MeanCorr_PS_= 0.65±0.04, p≤0.001). Fig. 2B (Table S4) illustrates the average correlation between each ROI and all other regions, indicating the regional nonspecific reduction of functional connectivity in the PS group. Fig. 2C represents the subtraction of the correlation matrices of the two groups displayed in Fig. 2A, which shows 71% (122 out of 171) of correlation comparisons are significantly different.

**Fig 2.**
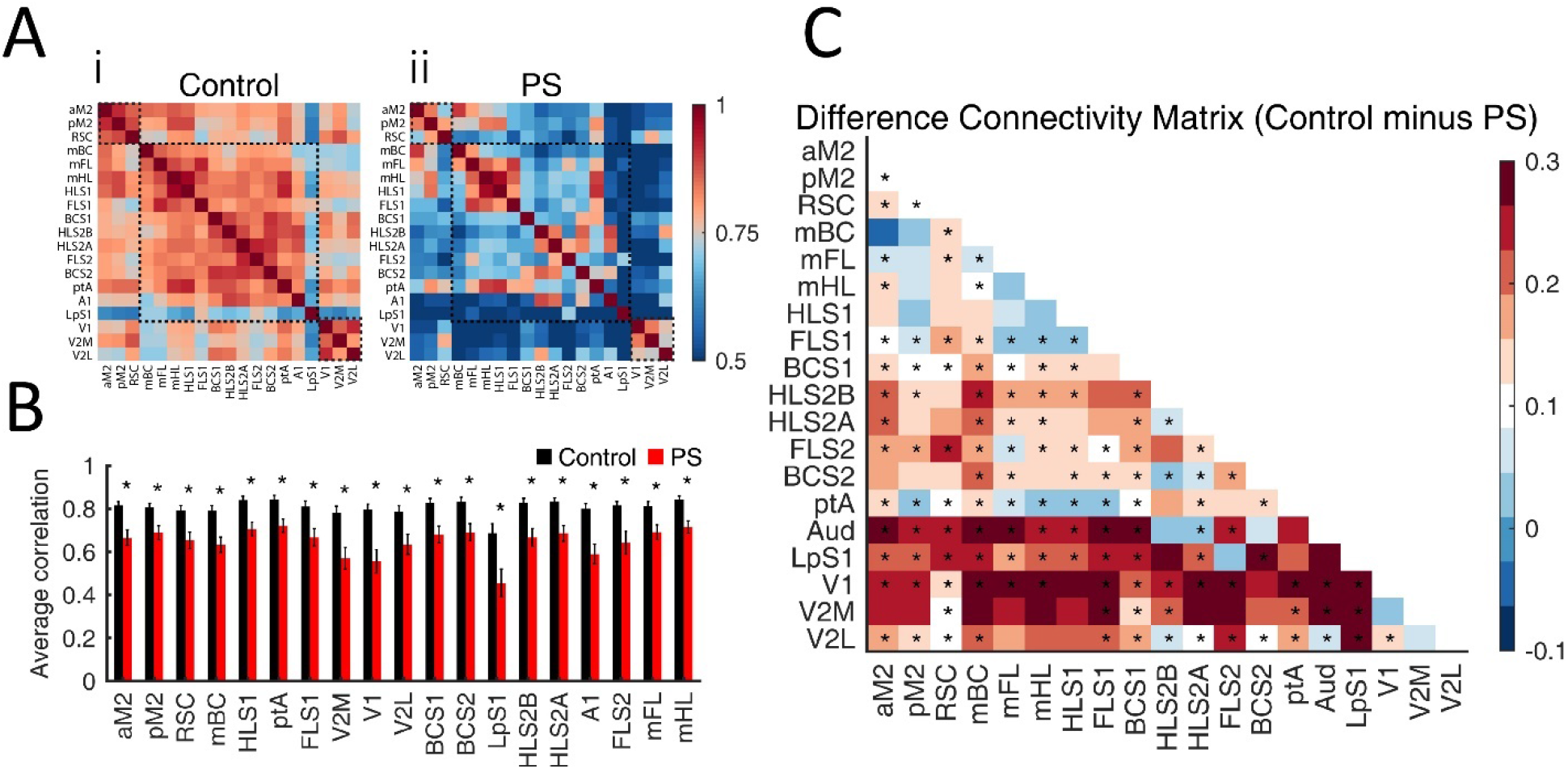
Reduced cortical functional connectivity by PS. **A**) Zero-lag correlation matrices for (i) the control and (ii) PS groups. Black dashed line squares separate midline, somatomotor, and visual regions (see the Methods and Fig. 1 for cortical regions). **B**) A significant difference was found between the two groups on the average correlation of each ROI and all other regions. Table S1 includes detailed statistics. P-values are corrected for multiple hypotheses testing. **C**) Subtraction of inter-regional correlation coefficients. The colors specify the degree of change in correlations. Correlation between most ROIs (71% (122 out of 171) was reduced in the PS. P-values are corrected for multiple hypotheses testing. The number of mice: n=12 control, n=6 PS. Asterisks indicate *p<0.05, **p<0.01, or ***p<0.001. Error bars indicate mean ± SEM (standard error of the mean). See the ROI identification in the Methods for the full name of ROIs.

### 3.4. Reduced efficiency and enhanced modularity in the PS network

Comparison of connectivity matrices in Fig. 2C shows a PS-induced overall decrease of correlation coefficient values within most of the regions compared with the control group. For a better understanding of intracortical relationships in the two groups, the correlation matrices were used to derive the corresponding graphs. The correlations were checked for statistical significance and no connection was removed as all p-values were much smaller than the significance level (see the Correlation and network analysis in the Methods). We did not apply proportion thresholding because as was explained in the Methods, all correlations were statistically significant at extremely high confidence levels, the size of the networks was small (19 nodes, 171 links), our findings were robust over a range of low thresholds, and high thresholds led to sparse small networks with poor statistics such that group differences were washed away. In Fig. 3A, graphs are plotted based on the average correlation in each group. The thickness of the connecting lines is proportional to the strength of the connection, and the nodes (regions) are positioned based on the average brain coordinates for animals. Both graphs were entirely connected, however, the links were thicker (i.e., larger correlation coefficients) in the control network.

**Fig 3.**
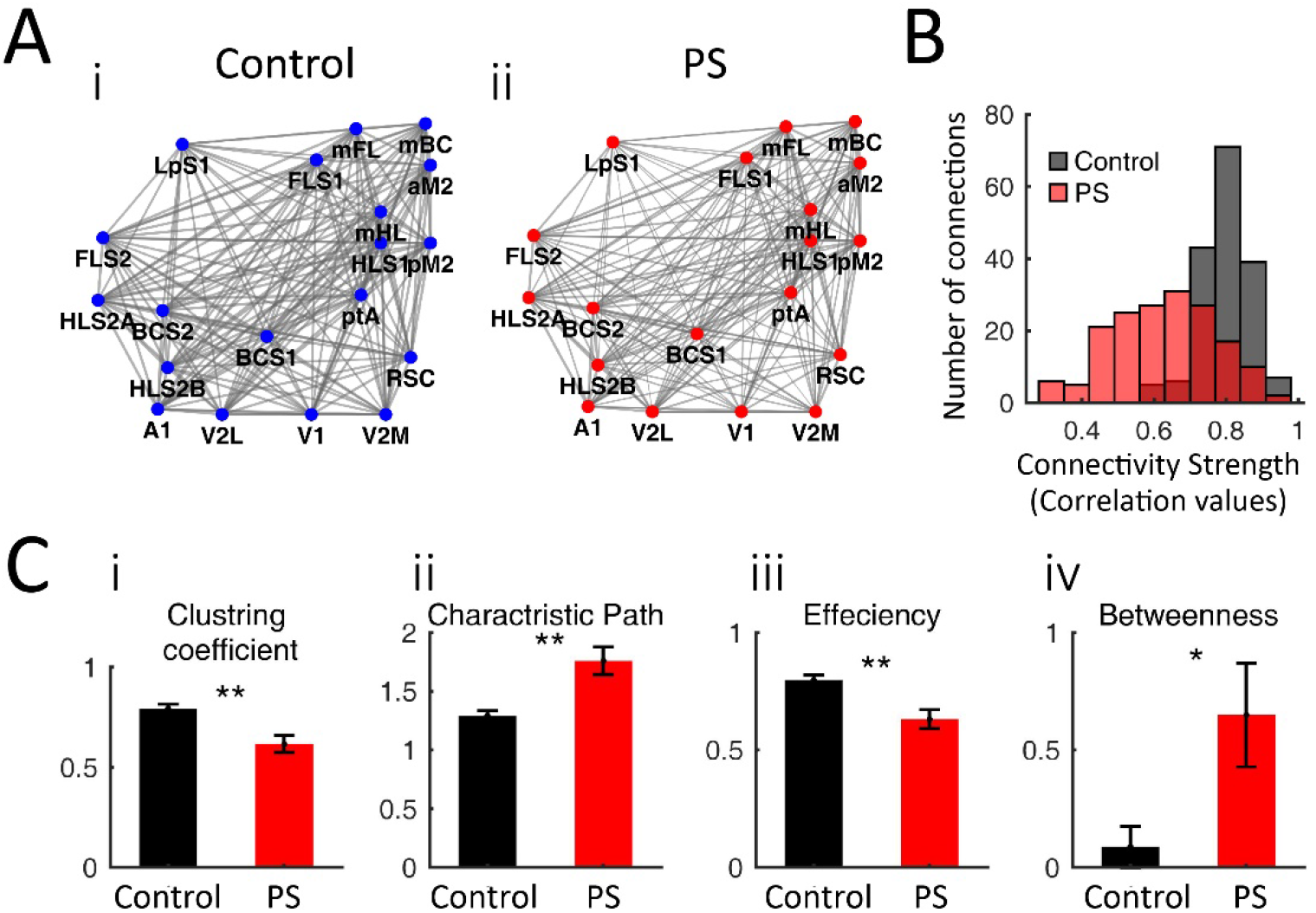
Declined efficient information transmission in the cortical network in PS. **A**) Schematic of the cortical networks based on the average correlation for (i) the control and (ii) PS groups. Nodes indicate cortical regions in the left hemisphere and edges’ weight represents the strength of the connection. Edges are thicker in the control network compared to the PS network, showing the existence of stronger connections among regions. **B**) The edge weight distributions are significantly different (Z=0.65 and p=1.2×10^-30^, two-sample Kolmogorov-Smirnov test). While edge-weights in the PS network spanned all values in 0 to 1 interval with a peak around 0.62, they only varied from 0.6 to 1 with a mean around 0.8 in the control group. **C**) Comparison of some global network measures. (i) The PS network showed a smaller clustering coefficient (df=16, sum of ranks=143, p-corrected =0.009), (ii) an increased characteristic path (df=16, sum of ranks=86, p-corrected=0.006), (iii) a lower global efficiency (df=16, sum of ranks=143, p-corrected=0.004) compared with the control network. (iv) The average betweenness of nodes was enhanced in the PS versus the control network (df=16, sum of ranks=93.5, p-corrected=0.035). P-values are corrected for multiple hypotheses testing. The bar graphs show the average value of the measures over all animals in a group. The number of mice: n=12 control, n=6 PS. Asterisks indicate *p<0.05, **p<0.01, or ***p<0.001. Error bars indicate mean ± SEM (standard error of the mean). See the ROI identification in the Methods for the full name of ROIs.

To find out how information flow might be affected by PS, we made a comparison of some basic global network measures indicating network efficiency such as “clustering coefficient”, “characteristic path length” and “global efficiency”. The “clustering coefficient” and the “characteristic path” are extensively used in brain network studies (Bullmore and Sporns, 2009). It has been shown that brain networks are highly clustered with a short characteristic path, which is a measure of “small-world property” (Hilgetag et al., 2000; Sporns and Zwi, 2004). Fig. 3C demonstrates a smaller clustering coefficient (Fig. 3Ci) and a longer characteristic path (Fig. 3Cii) of the PS network, which refer to a compromised small-world property in the PS mice. The “global efficiency” was also significantly reduced in the PS network (Fig. 3Ciii) which agrees with the increased characteristic path, as they are inversely related.

Using “betweenness centrality” as a measure of a region’s importance, we found that the average betweenness of nodes was increased in the PS compared to the control network (Fig. 3Civ). However, None of the ROIs consistently emerged as a central region among most animals in either of the groups.

To verify that the above group differences in network measures stemmed from the overall difference in connectivity strength distributions, Fig. 3B, we randomly permuted correlation matrices entries (see the Correlation and network analysis in the Methods). Under shuffling, group differences remained significant while there was no significant difference in the network measures within a given group. This confirms that the origin of all group differences in network charactristics is encoded in the weight distributions. See Supplementary Table S5 for the statistics.

In the next step, to detect segregated structure and hubs, two major hallmarks of brain networks (Bullmore and Sporns, 2009), partial correlation was used to capture the direct functional connection between nodes by removing confounding effects of third-party nodes (Smith et al., 2011; Smith et al., 2013) (see Supplementary Fig. 4 for partial correlation matrices). To limit the number of spurious connections from the estimating process of partial correlation (Martin et al., 2017; Oliver et al., 2019), all partial correlations were tested for statistical significance (Epskamp and Fried, 2018). Then the strongest 80% of connections in each network were preserved to equalizes the network sizes. As the networks are small (19 nods and 171 possible connections), more reduction of the links would be unreliable due to statistical fluctuations (see the Correlation and network analysis in the Methods for the details). Weighted networks in Fig. 4A display functional connectivity based on average partial correlation in each group. The edge widths imply the strength of the connection between regions and nodes with the same color belong to the same community. The negatively weighted connections in the partial correlation networks were also preserved as they could be neurobiologically associated with stress (see the Correlation and network analysis in the Methods). Positively and negatively weighted connections are respectively presented as green and red edges in Fig. 4A.

**Fig 4.**
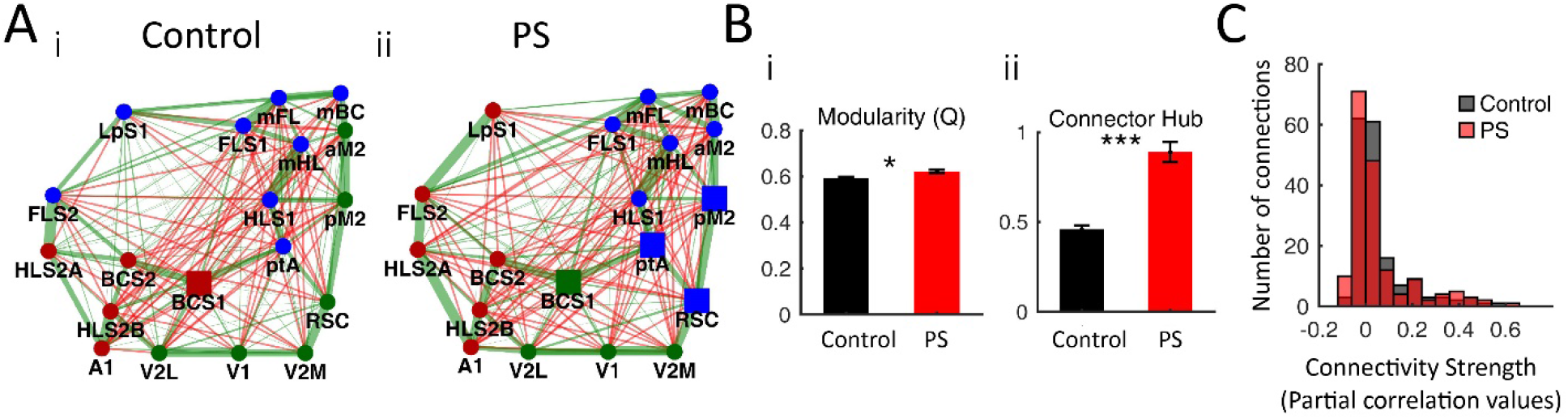
PS results in delineated modules in the cortical networks. **A**) Schematic of the cortical network based on the average partial correlation for (i) the control and (ii) PS networks. Nodes show cortical regions in the left hemisphere and edges’ weight reflects the strength of the connection. Green and red edges show positively and negatively weighted connections, respectively. The same color nodes belong to the same community. Square represents connector hubs. **B**) Bar graphs compare network measures between the two groups. (i) The PS network showed a more modular structure as the average of its modularity quality function (Q) was greater than the control network (df=16, sum of ranks=87, p-corrected=0.014). (ii) The number of connector hubs, high degree nodes with the participation coefficient of greater than 0.3, increased in the PS network (df=16, sum of ranks=78, p-corrected=3.2×10^-4^). P-values are corrected for multiple hypotheses testing. The bar graphs show the average value of the measures over all animals in a group. **C**) The edge weight distributions are not significantly different (Z=0.11 and p=0.227, two-sample Kolmogorov-Smirnov test). The number of mice: n=12 control, n=6 PS. Asterisks indicate *p<0.05, **p<0.01, or ***p<0.001. Error bars indicate mean ± SEM (standard error of the mean). See the ROI identification in the Methods for the full name of ROIs.

A higher number of non-overlapping groups of nodes in a given network is an indication of optimal network structures, and modularity is a quantified measure that shows to what extent a network is divided into such specified communities (Newman, 2006). Using community detection algorithms (see the Correlation and network analysis in Methods), a consensus partition for networks of both groups was created (Fig. 4A). Whereas the number of communities remained the same in both networks, the host modules of some nodes were different between the two groups. To make a quantified comparison of modularity, the average quality functions (Q, obtained from the community detection algorithm) were compared. Higher modularity in the PS network (Fig. 4Bi) implied that intermodular communication was more clearly separated from intramodular communication than the control network.

Generally, hubs are characterized as high degree nodes with edges that pose them in central positions for promoting communication through a network. To identify hubs in partial correlation networks, the participation coefficient was employed (Power et al., 2013). Based on their participation coefficient, “connector hubs” were categorized as high degree nodes with many intermodular connections (Sporns et al., 2007) (see the Correlation and network analysis in Methods). Fig. 4Bii shows that the average number of hubs is almost twice in the PS group. Regions shown as large squares in the networks in Fig. 4A emerged as hubs in the majority of mice in each group (BCS1 in the control group and BCS1, ptA, RSC, and pM2 in the PS group).

Fig. 4C shows that in contrast to the correlation values (Fig. 3B), there is no drastic difference in the histograms of the partial correlation values at the group level, reflecting the effect of network structure on the measure differences rather than the effect of the overall difference in connectivity strength distributions.

### 3.5. Rate and temporal order of motifs

To examine how PS alters patterns of activity in the brain, the rate and succession of the spontaneous sensory motifs were compared between the two groups. These spontaneously occurring motifs were determined using a template matching method through the templates extracted from multimodal sensory-evoked responses, Fig. 5A and S5. Using this approach, the spontaneous motifs originated from the forelimb, hindlimb, barrel, visual and auditory cortices were identified (Fig. 5Aii). The heat map in Fig. 5B demonstrates the concurrent correlation of different templates of sensory-evoked responses with a 30s period of spontaneous activity. The correlation of instantaneous patterns of spontaneous activity with multiple templates of sensory-evoked responses for the first 6s of the data in panel B has been plotted in Fig. 5C. Peaks in this plot reflect parts of spontaneous activity closely resembling the sensory-evoked templates. Shaded rectangles in Fig. 5C are recognized as motifs, whose VSD montage plots are shown in Fig. 5Aii (see Methods). The overall occurrence rate (i.e., number of motifs per second) was 0.31±0.01 and 0.29±0.01 in the control and PS groups respectively (p>0.1). Despite an almost identical overall rate of motifs’ occurrence in both groups, the occurrence frequency was increased for auditory motifs and decreased for forelimb motifs, with no changes for the visual, hindlimb, and whisker motifs (Fig. 5D).

**Fig 5.**
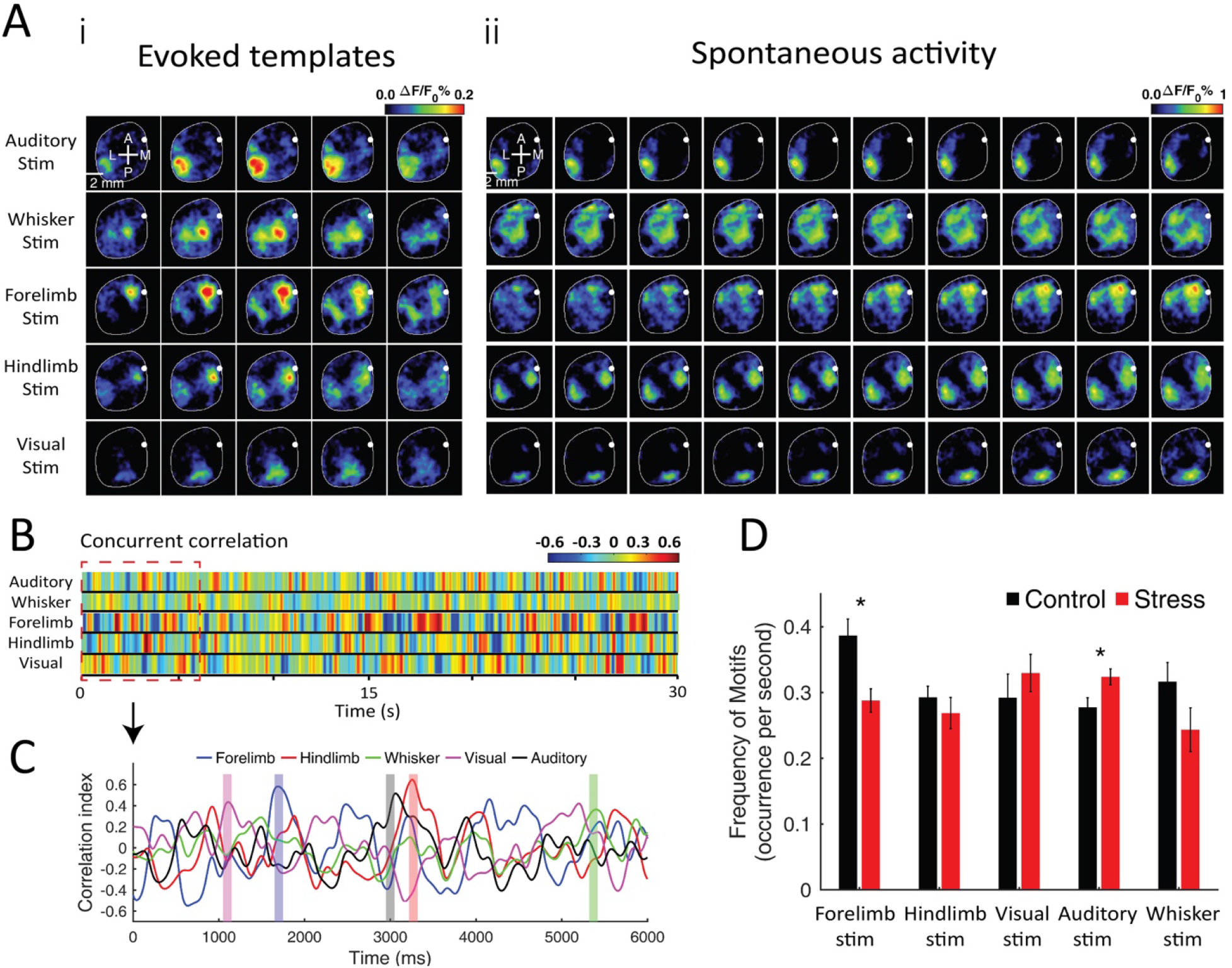
Increased frequency of auditory motifs and decreased frequency of forelimb motifs in PS. **A**) Comparison of sensory-evoked templates and spontaneous data from VSD imaging. (i) Five frames of the sensory-evoked responses to different stimuli. (ii) Spontaneous sensory motifs matched with evoked responses to auditory, whisker, forelimb, hindlimb, and visual stimuli. **B**) Concurrent correlation of different templates of sensory-evoked responses with a 30s period of spontaneous activity. The colors indicate the correlation value at different times. **C**) Correlation of sensory-evoked templates and spontaneous activity taken for 6s of spontaneous data (corresponding to the red dashed line in B). Shaded rectangles represent parts of spontaneous activity closely resembling the sensory-evoked templates, which were chosen as sensory motifs. **D**) The number of sensory motifs occurrence per second averaged over all animals in each group. The rate of occurrence in the PS group was decreased for the forelimb motif (df=12, sum of ranks=27, p=0.02) and increased for the auditory motifs (df=12, sum of ranks=61, p=0.04) compared with the control group. For other motifs no significant difference was observed (hindlimb: df=12, sum of ranks=36, p=0.27; visual: df=12, sum of ranks=51, p=0.5; whisker: df=12, sum of ranks=33, p=0.142). The number of mice: n=8 control, n=6 PS. Asterisks indicate *p<0.05, **p<0.01, or ***p<0.001. Error bars indicate mean ± SEM (standard error of the mean).

While spontaneous cortical activity has been shown to constitute a complex combination of multiple sensory motifs (Afrashteh et al.; Mohajerani et al., 2013), their sequential order remains unknown. Thus, we tested whether PS alters the temporal order of motif combinations by comparing all possible nonrepetitive permutations, including, 20 double, 60 triple, 120 quadruple, and 120 quintuple sequences (see Methods). The results of quintuple sequences, however, were not included given their rare occurrence. Besides comparing PS and control groups, the temporal order of motif sequences in each group was also compared with random data (see Methods).

Fig. 6 exhibits the average number of sequences that revealed a significant difference between experimental and random data in both groups (quadruple sequences: control, 0.088±0.015, control-shuffled, 0.140±0.006, p> 0.02; PS, 0.087±0.01, PS-shuffled, 0.13±0.006, p>0.008; triple sequences: control, 0.51±0.064, control-shuffled, 0.72±0.027, p>0.02; PS, 0.61±0.1, PS-shuffled, 0.67±0.028, p>0.015; double sequences: control, 3.05±0.22, control-shuffled, 3.7±0.12, p>0.04; PS, 2.92±0.18, PS-shuffled, 3.45±0.13, p>0.04). Despite significant change in the average rate of sequences, few individual sequences in each group emerged significantly different from the random occurrence (Supplementary Fig. 6A, 6B, and 6C). According to Fig. 6, PS also had no effect on the average rate of multiple combinations’ occurrence (quadruple sequences: control, 0.088±0.015, PS, 0.087±0.01, p>0.85; triple sequences: control, 0.51±0.064, PS, 0.5±0.045, p>0.66; double sequences: control, 3.05±0.22, PS, 2.92±0.18, p>0.75) (see Supplementary Fig. 6D for comparison between individual sequences).

**Fig 6.**
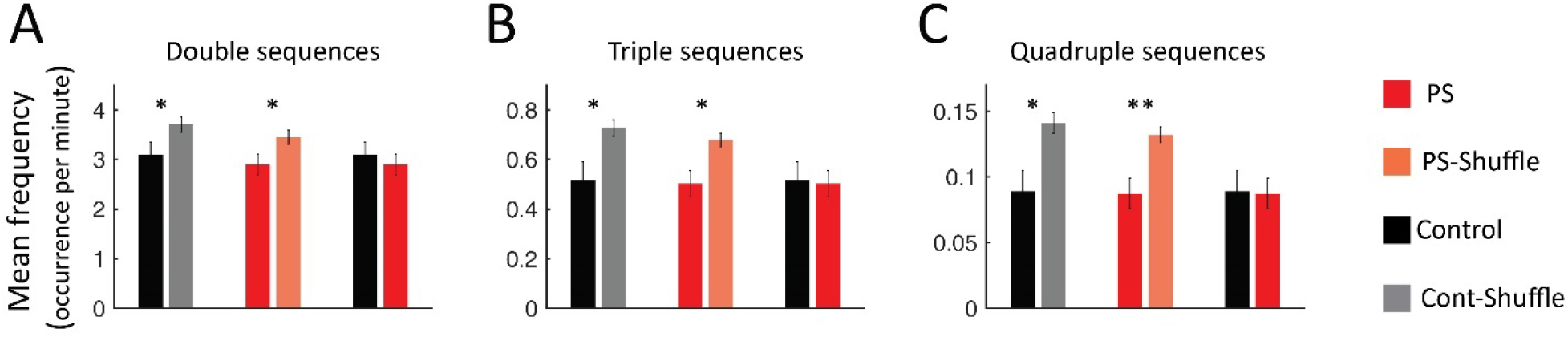
Nonrandomly occurrence of sensory motifs. Bar graphs indicate the average frequency of nonrepetitive **A**) double, **B**) triple, and **C**) quadruple sequences of sensory motifs. Whereas, the two groups were similar in the occurrence rate of motif sequences, they significantly differ from their corresponding shuffled double, triple, and quadruple combinations (Double: Cont-Cont-Shuffle, df=12, sum of ranks=49, p=0.05, Str-Str-Shuffle, df=12, sum of ranks=26, p=0.04; Triple: Cont-Cont-Shuffle, df=12, sum of ranks=45, p=0.012, Str-Str-Shuffle, df=12, sum of ranks=23, p=0.008; Quadruple: Cont-Cont-Shuffle, df=12, sum of ranks=46, p=0.02, Str-Str-Shuffle, df=12, sum of ranks=21, p=0.002;). The number of mice: n=8 control, n=6 PS. Asterisks indicate *p<0.05, **p<0.01, or ***p<0.001.

## 4. Discussion

The experiments examined how exposure to stress during the prenatal period affects the neocortical function of adult mice. Measures were made of cortical SERs, RSFC, and spontaneous sensory motifs in PS mice compared with the control mice at 2-3 months of age. PS was associated with an elevated corticosterone level, the reduced amplitude of SERs, overall decline of RSFC and network efficiency, enhanced modularity, an increased number of connector hubs, and the altered occurrence frequency of auditory and forelimb motifs.

### 4.1. Elevated corticosterone levels

In our study, the PS mice showed a four-fold higher basal corticosterone level than the control group in adulthood. Corticosterone is the primary component of the stress system that along with the sympathetic nervous system contributes to mobilizing energy stores in response to stressful events (Charil et al., 2010; Handa and Weiser, 2014). Current experimental research supports the idea that the dysregulation of the HPA-axis during sensitive fetal periods is associated with lasting alterations in brain and behavior throughout the lifespan (Charil et al., 2010; Huizink et al., 2004; Weinstock, 2017). Findings of studies also demonstrate that prenatal noise stress alone or in combination with other stresses is linked to higher levels of corticosterone at rest, delay in sensory and motor development, and life-long impairments in behavioral and cognitive performance (Jafari et al., 2020b).

Adrenal glucocorticoid hormones are essential for the maintenance of homeostasis and adaptation to stress. They act via the mineralocorticoid receptors (MRs) and glucocorticoid receptors (GRs) (Graham et al., 2019), which are co-expressed abundantly in the neurons of limbic structures, including the hippocampus, mPFC, and amygdala (Czeh et al., 2007; de Kloet et al., 2005; Kim et al., 2015). It has been shown that long-term exposure to stress (such as was indicated by a high basal level of corticosterone in our study) or corticosterone administration downregulates the glucocorticoid receptors and increases the responsivity of the HPA-axis to stress. This effect builds a vicious neuroendocrine circle of increased corticosterone levels and increased responsiveness to stress (Eggermont, 2014; Joels et al., 2006).

The limbic brain is the primary neural circuitry that mediates the response to stressful events (Czeh et al., 2007). The amygdala makes a connection between subcortical areas, which are involved in the fear response, and cerebral cortical regions, which receive sensory information from the external environment (Wilson et al., 2015). The auditory system connects to the autonomic nervous system (ANS) (including brain mechanisms underlying, stress, arousal, startle, and blood pressure) via the amygdala and other circuits (Burow et al., 2005; Eggermont, 2014; Jafari et al., 2020b). Thus, noise can activate non-classical auditory-responsive brain areas, trigger the emotion/fear system of the brain (Chen et al., 2016; Eggermont, 2014), and lead to the release of stress-related hormones, including corticotropin-releasing hormone (CRH), corticosterone, and norepinephrine (Burow et al., 2005; Eggermont, 2014). Considering the connection between the auditory system and the neural system underlying stress signaling, cortical auditory responses might show more susceptibility to noise stress than other sensory responses. Animal studies also indicate that enduring HPA dysregulation leads to dendritic atrophy, reduced neurogenesis, altered synaptic efficacy and receptor distribution, and impaired synaptic plasticity in the limbic system (Joels et al., 2004; McEwen, 2004), as well as reduced volume in the corpus callosum and frontal cortical areas (Amaya et al., 2021).

Human studies also indicate that prenatal stressors during sensitive periods of development have an organizational effect on biological systems, and systematically pose a potential risk factor for developing multiple neuropsychiatric disorders during the lifespan (Abbott et al., 2018; Van den Bergh et al., 2017). They can result in epigenetic variations of DNA (deoxyribonucleic acid)-methylation at CRH and glucocorticoid receptor (GR) gene promoters lasting into adulthood and can even continue to the next generation (Bock et al., 2015; Crudo et al., 2012). In a recent study on young adults, higher prenatal stress was related to more mood dysregulation, lower overall gray matter (GM) volume, and lower GM volume in the mid-dorsolateral frontal cortex, anterior cingulate cortex, and precuneus (Marecková et al., 2019). Findings of current studies also show the relationship between maternal cortisol during pregnancy and resting-state functional connectivity in human infants (Graham et al., 2019; Scheinost et al., 2020).

### 4.2. Decreased cortical evoked responses to sensory stimuli

In this study, the amplitude of evoked responses to sensory stimuli was significantly reduced for the PS mice relative to the control group. Our findings were consistent with past animal studies that indicate the adverse effect of increased corticosterone levels on SERs. For instance, it has been shown that corticosterone administration during the first two postnatal weeks is associated with delayed development of cortical evoked responses to auditory, visual, and sciatic nerve stimulation (Salas and Schapiro, 1970) and the startle reflex (Pavlovska-Teglia et al., 1995). In another study on adult C57BL/6NJ mice, changes in the amplitude of auditory evoked responses due to chronic corticosterone administration were dose-dependent, and the serum corticosterone concentration was negatively correlated with the amplitude of auditory evoked responses (Maxwell et al., 2006). The amplitude of forelimb stimulation also was reduced in a study on adult C57BL/6NJ mice under chronic restraint stress (Han et al., 2019). Findings of other studies also point to reduced sensory gating subsequent to chronic corticosterone administration, which may lead to abnormalities in sensory information processing (Maxwell et al., 2006; Stevens et al., 2001). It has been suggested that the suppress of sensory information due to increased corticosterone level may result from an adaptive response to immediate unexpected stimuli (Devilbiss et al., 2012), which should be further studied in future research.

### 4.3. Reduced efficiency of the cortical network

In this study, the overall functional connectivity (i.e., correlation strength), regional cortical connectivity (i.e., the average correlation between one brain ROI and all other regions), and network efficiency were declined in the PS mice relative to the control group. We applied several network measures of brain connectivity (e.g., clustering coefficient, characteristic path length, global efficiency) that were all in support of reduced efficiency of information flow (Rubinov and Sporns, 2010) in the PS network

Evidences from human brain imaging techniques demonstrates long-lasting PS-associated structural and functional changes in several brain regions (e.g., the prefrontal, parietal, and temporal lobes, and the cerebellum, hippocampus, and amygdala) (van den Bergh et al., 2018). Alterations in brain functional connectivity have been shown in amygdalar–thalamic networks and intrinsic brain networks (e.g., DMNs and attentional networks) (Favaro et al., 2015; Graham et al., 2016; Scheinost et al., 2016; Turesky et al., 2019). Animal studies also indicate that PS or early exposure to environmental stresses alters synaptogenesis in the neocortex and the hippocampus (Bale et al., 2010; Mychasiuk et al., 2012), impacts brain development (Kolb et al., 2013; Mychasiuk et al., 2012; Weinstock, 2017), and produces alterations in the connectome (Scheinost et al., 2017). In a recent study in male offspring of pregnant C57BL/6NG mice exposed to restraint stress during the last week of pregnancy, the PS was associated with the reduction of temporal coupling between neuronal discharge in the medial prefrontal cortex (mPFC) and hippocampal sharp-wave ripples (Negron-Oyarzo et al., 2015).

It has been shown that the PS-induced dysregulation of the HPA-axis drives epigenetic changes in the developing brain (Scheinost et al., 2017). Cortisol (i.e., corticosterone in animals), the end product of the HPA-axis, has a prominent contribution to brain homeostasis and the response to various types of psychological and environmental stresses (Sapolsky et al., 2000). Increased maternal cortisol levels during pregnancy can influence the fetal brain through the placenta or stimulating fetal cortisol production (Sapolsky et al., 2000). The placental enzyme 11β-HSD2, which is a partial barrier to the passage of active cortisol, is downregulated in adverse prenatal conditions (Reynolds, 2013). In two recent studies, the association of maternal cortisol concentrations during pregnancy and the offspring’s brain functional connectivity was sex-specific (Graham et al., 2019; Kim et al., 2017). For instance, elevated maternal cortisol was associated with reduced amygdala connectivity to several brain regions involved in sensory processing and integration, DMN, and emotion perception in boys, and stronger connectivity to these brain regions in girls (Graham et al., 2019). This finding might be associated with the etiology of gender differences in internalizing psychiatric disorders (Liu et al., 2011).

Both human and nonhuman studies point to altered maternal immune activity in PS, which poses offspring at risk for cognitive and behavioral disorders (Estes and McAllister, 2016; Scheinost et al., 2017). In a study in human neonates at 40–44 weeks postmenstrual age, maternal third trimester interleukin-6 (IL-6) and C-reactive protein levels, which regulate immune responses, were related to brain functional connectivity in default mode, salience, and frontoparietal networks, as well as with both fetal and toddler behavior (Spann et al., 2018). IL-6, which crosses both the placenta and blood-brain barrier (Zaretsky et al., 2004), is part of both the pro-inflammatory and anti-inflammatory pathways and their imbalance contributes to abnormal cognitive and behavioral development (Meyer et al., 2008). It also has been found that early life experiences and the resulting behavioral consequences can be transmitted to the next generation through the epigenetic processes, which indicates a transgenerational cycle of changes in the brain and behavior (Bock et al., 2014). Together, several mechanisms, including alterations in the neuroendocrine system, the immune system, and the epigenetic processes may be implicated in reduced efficiency of information flow and undermined small-world property (Rubinov and Sporns, 2010) in PS mice, which suggest directions for future studies.

### 4.4. Increased cortical network modularity

To further investigate the impact of PS on brain network connectivity, “modularity”, the degree a network can be classified into non-overlapping but internally interactive subsets of regions (Thomason et al., 2014), and hubs were examined. PS network demonstrated enhanced modularity (i.e., more disjointed but tight modules) and increased average number of connector hubs (i.e., high degree nodes contributing widely among communities (Sporns et al., 2007)) relative to the control network.

Whereas few studies have shown how network modularity and nodes’ connections are changing in rodent brains in the context of PS, it has been found that hubs are biologically costly, because they have higher metabolic demands and longer-distance connections compared to other brain regions (Crossley et al., 2014). Studies also suggest that the integrative role of hub nodes makes them more vulnerable to dysfunction in brain disorders (van den Bergh et al., 2018), in which deviant hubs may show more tendency to function in brain pathological states (Crossley et al., 2014; Liska et al., 2015; Sporns, 2014). Thus, our findings might be interpreted as showing that PS is associated with enhanced contribution of primary barrel cortex (BCS1), parietal association (ptA), posterior secondary motor (pM2), and retrosplenial (RSC) cortices in communication between networks and response to environmental stimuli. It might be inferred that increased modularity in the PS network is an indication of more segregated functional sub-networks that were not ameliorated with increasing age.

### 4.5. Altered frequency of spontaneous sensory motifs and their temporal order

Multiple anatomically consistent resting-state networks in the brain (e.g., DMNs, sensorimotor, and cortical sensory areas) produce temporally coherent spontaneous fluctuations at rest or in the absence of any external stimulation (Kong et al., 2014). A group of these intrinsic responses, which originate from primary sensory cortices, expand identical to SERs called sensory motifs (Kenet et al., 2003). It has been found that the brain does computations using sensory motifs (Turkheimer et al., 2015). Among five spontaneous sensory motifs (e.g., visual, auditory, whisker, forelimb, and hindlimb) examined in this study, the PS was associated with an increase in the frequency of auditory motifs and a decrease in the rate of forelimb motifs. In the McGirr et al. study (McGirr et al., 2020) using a postnatal maternal deprivation stress paradigm over whisker, forelimb, and hindlimb sensory motifs, an increased occurrence of whisker and forelimb motifs was reported. These findings indicate that stress does not uniformly alter the frequency of different sensory motifs, which may result from the stress procedure applied (e.g., time, duration, and type of stress exposure). Therefore, enhanced auditory motif’s frequency in our study might be due to the applied auditory stress paradigm making auditory motifs more susceptible to stress (refer to section 4.1, paragraph 3). The reduction of forelimb motifs, however, is unclear since little evidence was shown in this area. Neurophysiological evidence also refers to the role of spontaneous sensory motifs in modulating cortical responses to sensory inputs (Ferezou and Deneux, 2017), as well as higher-level cognitive operations and behavior (Turkheimer et al., 2015). In light of these findings, the stress-related alterations of sensory motifs and their contribution to sensory processing and psychiatric disorders should be further examined in the future.

Studies using different recording techniques have shown that cortical spontaneous activity is a non-random process that represents the relationship among neurons (Azouz and Gray, 1999), cortical columns (Smith et al., 2018), and neuroanatomical systems (Xiao et al., 2017). In addition, from the network analysis perspective, the cortex has been ubiquitously reported as having a non-random organization (Sporns, 2011). In our study, the sequential order of sensory motifs in spontaneous neural activity was considered as a measure of randomness. In agreement with the experimental evidence of nonrandom cortical activity (Azouz and Gray, 1999; Smith et al., 2018; Xiao et al., 2017), the sequences of sensory motifs occurred in a nonrandom order in all animals in our study, and no PS effect was observed.

## Conclusions

This study aimed to investigate how PS alters sensory-evoked responses (SERs), resting-state functional connectivity (RSFC), and the occurrence of sensory motifs in the adult mouse brain. The PS was associated with reduced amplitude of all cortical SERs, an overall decrease of RSFC and network efficiency, enhanced modularity or segregated functional sub-networks, and altered occurrence frequency of auditory and forelimb motifs. Four connector hubs were also found in the PS network that may function as brain regions with higher vulnerability to dysfunction. Future studies are suggested to investigate: 1) the association of altered cortical SERs, intrinsic FC, spontaneous sensory motifs, and identified hubs with PS-related behavioral disorders; and, 2) the impact of postnatal pharmacological or behavioral treatments in reversing or alleviating the PS impacts. The present study was performed under the isoflurane-anesthetized state in male mice. The replication of our experiments in both sexes during awake and anesthetized states will help to understand how the results are influenced by brain states and sex.

## Acknowledgments

The present review was supported by Canadian Institutes of Health Research (CIHR) Grant# 390930, Natural Sciences and Engineering Research Council of Canada (NSERC) Discovery Grant #40352, Alberta Innovates (CAIP Chair) Grant #43568, Alberta Alzheimer Research Program Grant # PAZ15010 and PAZ17010, and Alzheimer Society of Canada Grant# 43674 to MHM. This study was part of a postdoctoral fellowship to ZJ and a Ph.D. degree to ZR in the Canadian Center for Behavioral Neuroscience (CCBN) at the University of Lethbridge. We would like to thank Javad Karimi Abadchi for his useful comments.

## Author Contribution

Conceptualization, M.H.M., Z.J., B.E.K.; Experimentation, M.H.M, Z.J, N.A.; Formal Analysis, Z.R.; Analysis advice, N.A., R.T., S.S., J.D.; Writing - Original Draft, Z.R., Z.J., M.H.M.; Writing - Review & Editing, Z.R., Z.J., B.E.K., J.D., and M.H.M. Funding Acquisition, M.H.M.; Resources, M.H.M.; Supervision, M.H.M.

## 5. Supplemental Information

**Supplementary Fig 1.**
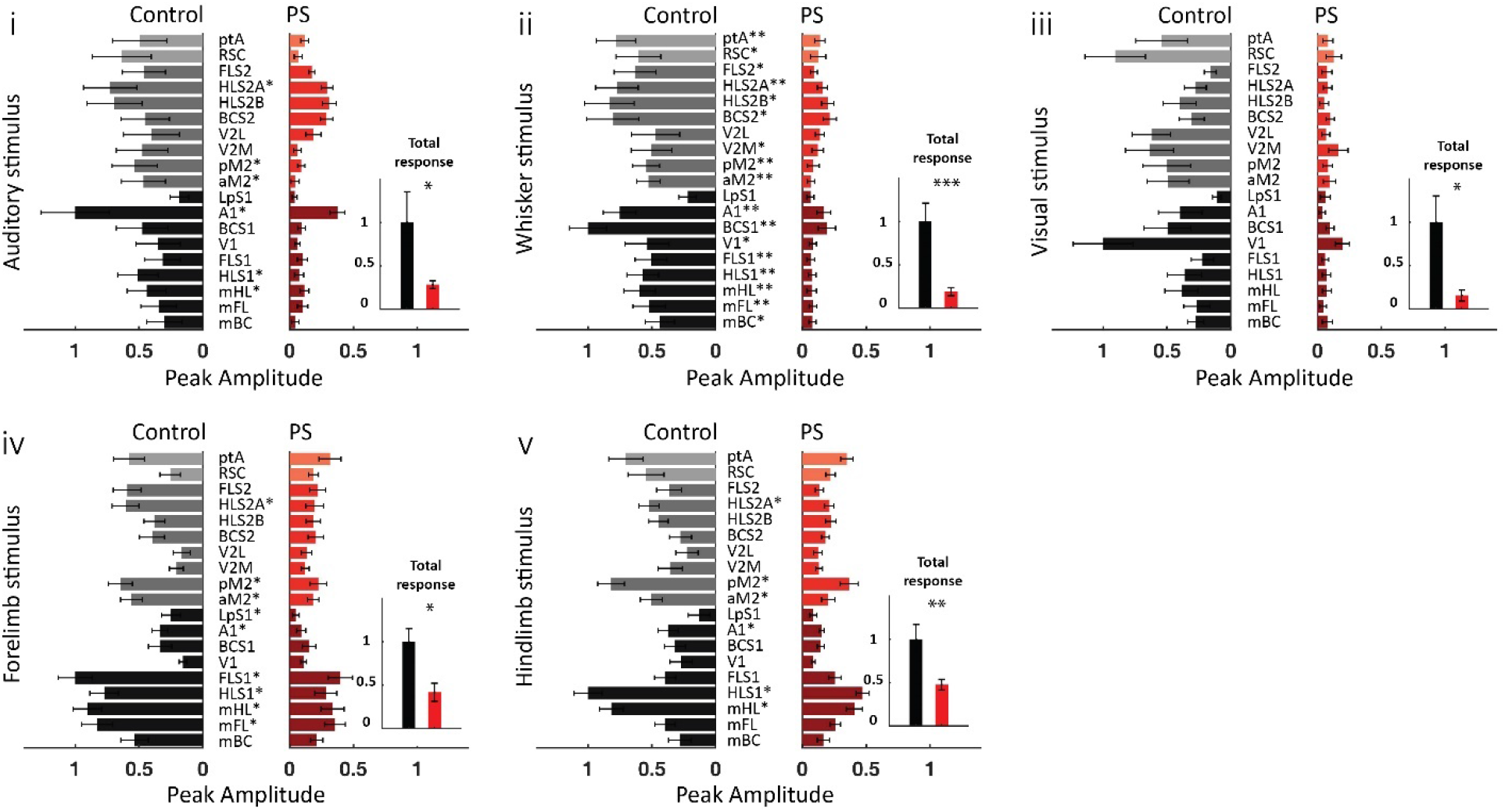
The reduced peak amplitude of evoked responses to different sensory stimuli in PS. Peak amplitude of response to (i) auditory, (ii) whisker, (iii) visual, (iv) forelimb, and (v) hindlimb stimuli. For each stimulation, bars represent the response of each region averaged over animals per group. Note that for each stimulation, all responses (from both control and PS groups) were normalized to the largest response, which always belonged to the primary region corresponding to that sensory stimulus in the control group. Cortical regions were divided into three groups of primary sensory and motor areas (mBC, mFL, mHL, HLS1, FLS1, V1, BCS1, A1, and LPS1), supplementary areas (aM2, pMa, V2M, V2L, BCS2, HLS2A, HLS2B, FLS2), and associative areas (ptA and RSC). The gray and red spectrum colors were used for the control and PS groups, respectively. Three categories of colors in each group from dark to light indicate primary, secondary, and associative areas, respectively. Table S3 provides related statistics. P-values are corrected for multiple hypotheses testing. Small inset bar graphs also compare the sum of the peak amplitudes of all regions, normalized to the control group’s values (auditory stimulus: df=13, p=0.017, sum of ranks=92; whisker stimulus: df=13, p=8×10^-4^, sum of ranks=98; forelimb stimulus: df=13, p=0.017, sum of ranks=92; hindlimb: df=13, p=0.007, sum of ranks=94; visual: df=13, p=0.036, sum of ranks=90). The number of mice: n=9 control, n=6 PS. Asterisks indicate *p<0.05, **p<0.01 or ***p<0.001. See the ROI identification in the Methods for the full name of ROIs.

**Supplementary Fig 2.**
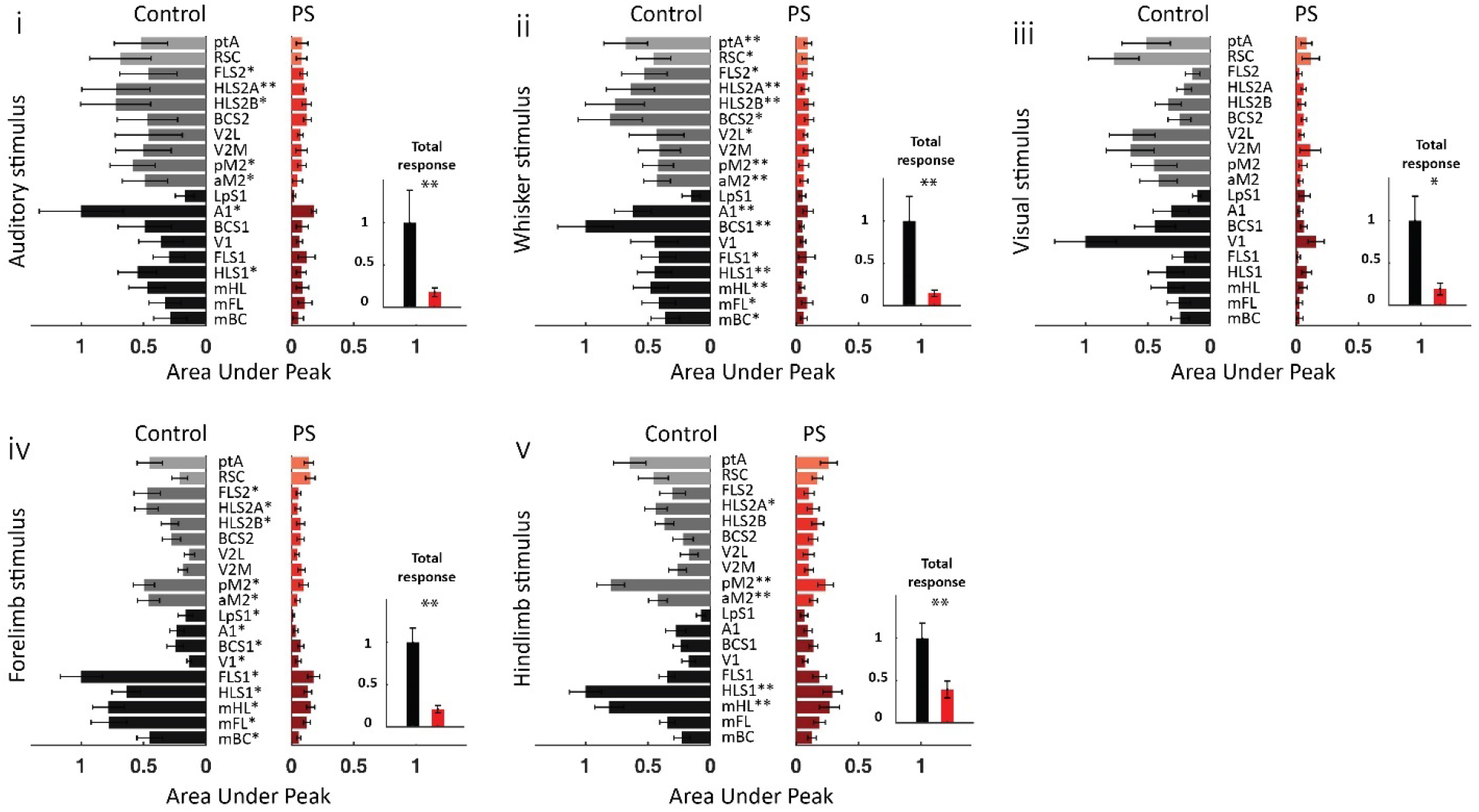
The reduced area under the peak of the evoked responses to different sensory stimuli in PS. Area under the peak of response to (i) auditory, (ii) whisker, (iii) visual, (iv) forelimb, and (v) hindlimb stimuli. For each stimulation, bars represent areas under the peak of the response of each region averaged over animals per group. Note that for each stimulation, all values (from both control and PS groups) were normalized to the largest one, which always belonged to the primary region corresponding to that sensory stimulus in the control group. Cortical regions were divided into three groups of primary sensory and motor areas (mBC, mFL, mHL, HLS1, FLS1, V1, BCS1, A1, and LPS1), supplementary areas (aM2, pMa, V2M, V2L, BCS2, HLS2A, HLS2B, FLS2), and associative areas (ptA and RSC). The gray and red spectrum colors were used for the control and PS groups, respectively. Three categories of colors in each group from dark to light indicate primary, secondary and associative areas, respectively. Table S4 includes related statistics. P-values are corrected for multiple hypotheses testing. Small inset bar graphs, also, compare sum of the area under peak of responses of all regions, normalized to the control group’s values (auditory stimulus: df=13, p=0.002, sum of ranks=96; whisker stimulus: df=13, p=0.004, sum of ranks=95; forelimb stimulus: df=13, p=0.007, sum of ranks=94; hindlimb: df=13, p=0.004, sum of ranks=95; visual: df=13, p=0.017, sum of ranks=92). The number of mice: n=9 control, n=6 PS. Asterisks indicate *p<0.05, **p<0.01 or ***p<0.001. See the ROI identification in the Methods for the full name of ROIs.

**Supplementary Fig 3.**
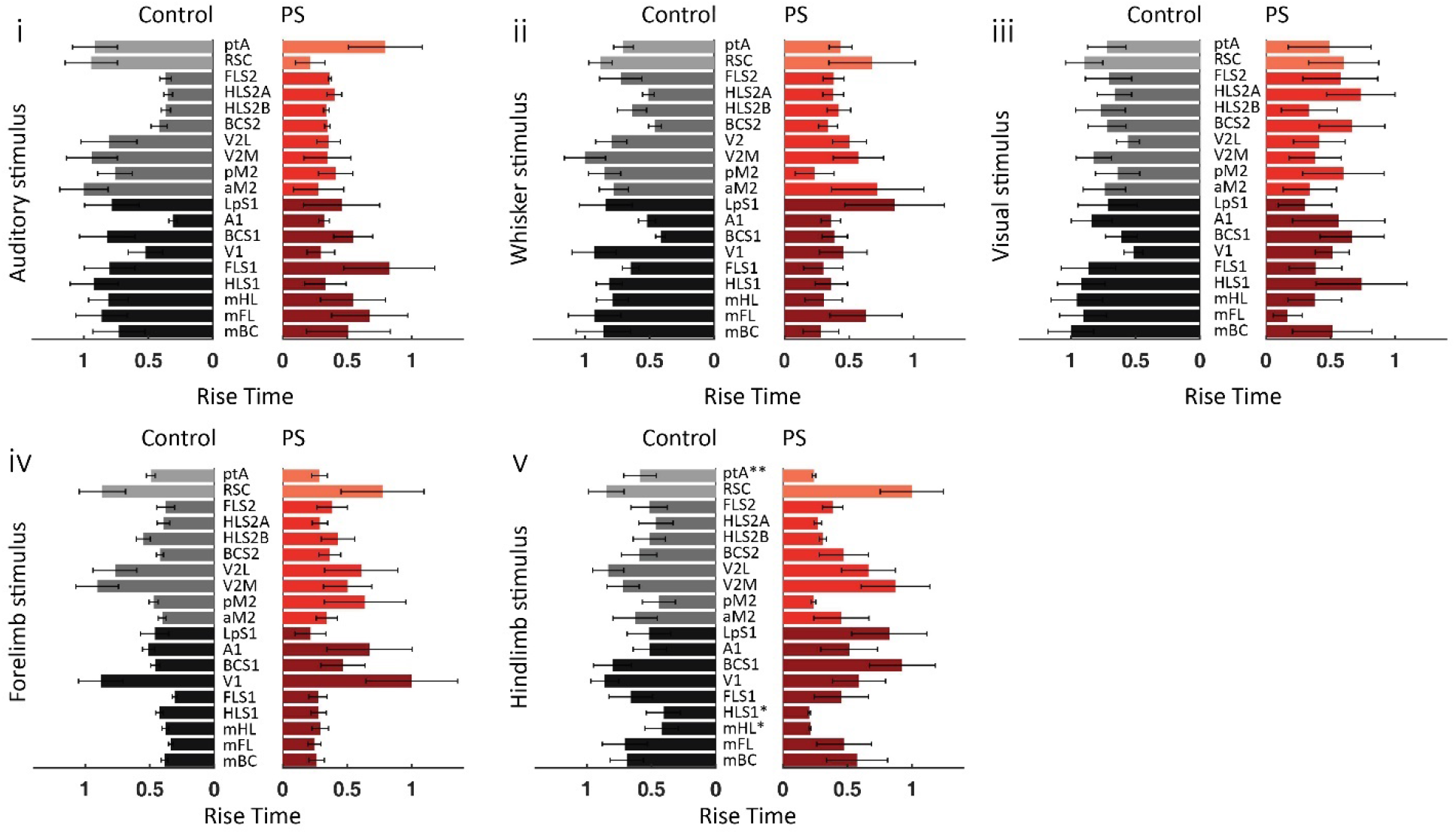
PS did not significantly affect the rise time of the peak of response to different sensory stimuli. Rise time of response to (i) auditory, (ii) whisker, (iii) visual, (iv) forelimb, and (v) hindlimb stimuli. For each stimulation, bars represent the rise time of response of each ROI averaged over animals per group. Note that for each stimulation, all values (from both control and PS groups) were normalized to the largest rise time. Cortical regions were divided into three groups of primary sensory and motor areas (mBC, mFL, mHL, HLS1, FLS1, V1, BCS1, A1, and LPS1), supplementary areas (aM2, pMa, V2M, V2L, BCS2, HLS2A, HLS2B, FLS2), and associative areas (ptA and RSC). The gray and red spectrum colors were used for the control and PS groups, respectively. Three categories of colors in each group from dark to light indicate primary, secondary and associative areas, respectively. None of the areas showed a significant difference in the rise time. Table S3 includes associated statistics. The number of mice: n=9 control, n=6 PS. See the ROI identification in the Methods for the full name of ROIs.

**Supplementary Fig 4.**
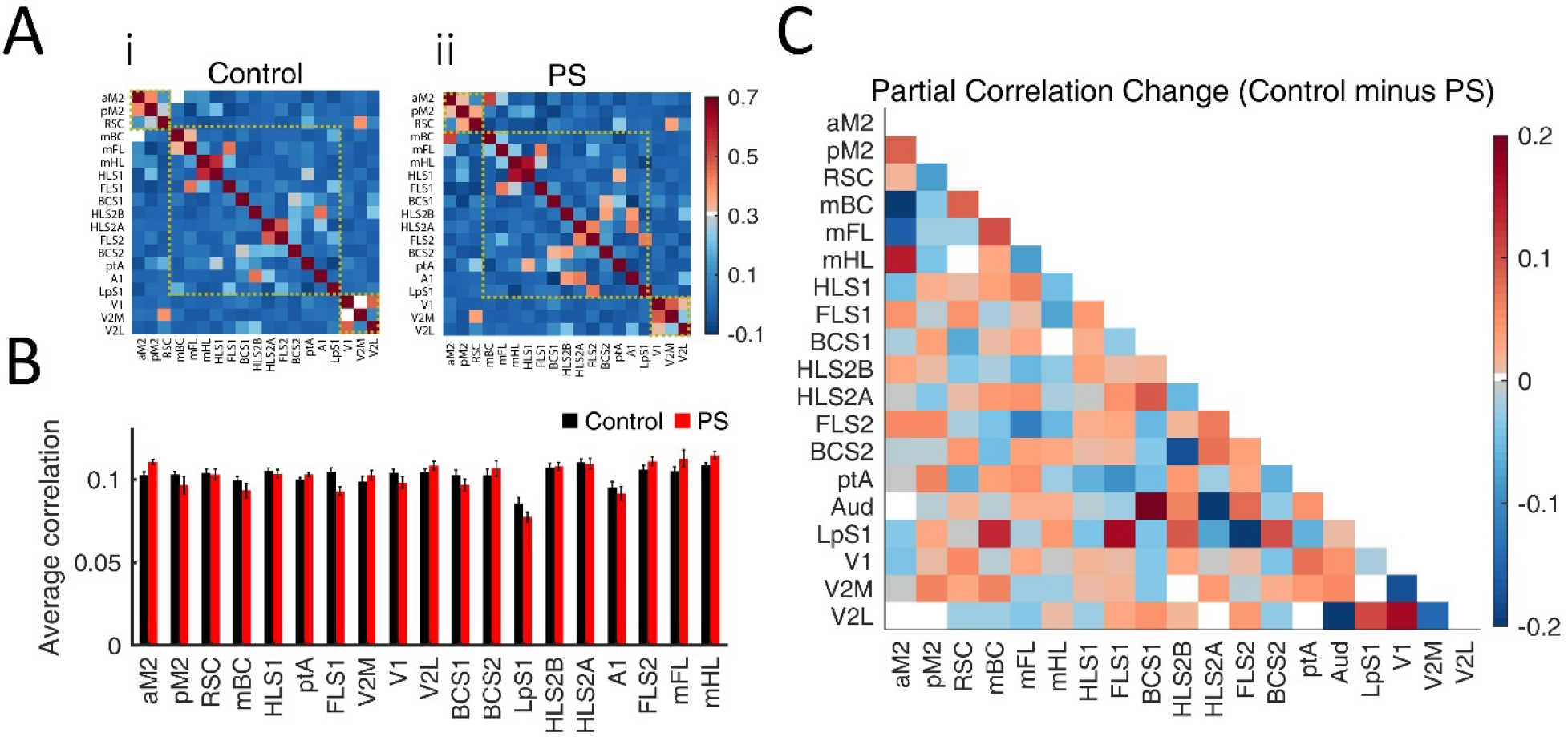
PS did not significantly change the partial correlation between cortical ROIs. **A**) Partial correlation matrices for (i) control and (ii) PS groups. Dashed line squares separate midline, somatomotor, and visual regions (see the Methods and Fig. 1 for cortical regions). **B**) The average of the partial correlation of every single region with all other regions did not show a significant difference between PS and control groups, after correcting for multiple hypotheses testing. **C**) Subtraction of inter-regional partial correlation coefficients. The colors identify the degree of change in correlation. Whereas the inter-regional partial correlation of several areas was decreased in PS, it was increased for other areas. The alterations, however, were not significant. The number of mice: n=12 control, n=6 PS. See the ROI identification in the Methods for the full name of ROIs.

**Supplementary Fig 5.**
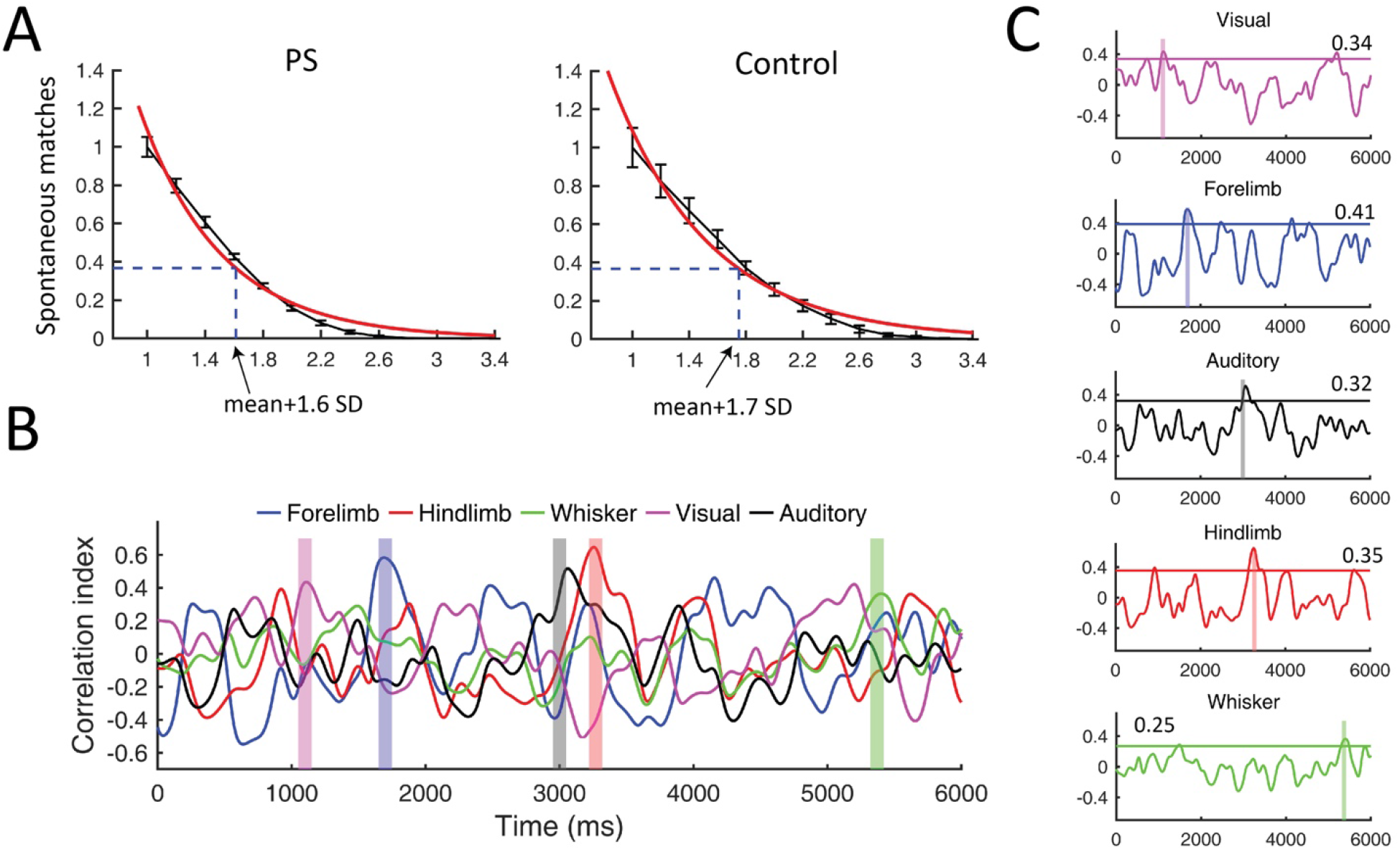
Relationship between the percentage of matched spontaneous sensory motifs and the threshold level. **A**) The mean correlation between each sensory template and spontaneous activity was summed with a standard deviation (SD) in steps of 0.1 SD to make a range of thresholds. For the forelimb sensory template, the number of ‘matches’ was plotted for the range of thresholds, which decreased exponentially. The threshold level was set where the percentage of matches reached 36% (e^-1^) of its initial value. To find the e^-1^ decay, the data (black plot) was fitted with an exponential function (red plot). The intersection of the dashed line and x-axis indicates the level of threshold. Although this figure shows the threshold level for finding forelimb sensory motifs in each group, the threshold level for all other sensory motifs also fell almost around their mean correlation+1.6 SD for the PS group and mean correlation+1.7 SD for the control group. **B**) Correlation of sensory-evoked templates and spontaneous activity taken for 6s of spontaneous data. Shaded rectangles represent parts of spontaneous activity closely resembling the sensory-evoked templates, which were chosen as sensory motifs after meeting the threshold criterion explained in methods. **C**) The correlation plots in panel B were shown separately with their threshold levels. The threshold values were the mean correlation between each sensory template and spontaneous activity plus 1.6 SD. The number of mice: n=8 control, n=6 PS.

**Supplementary Fig 6.**
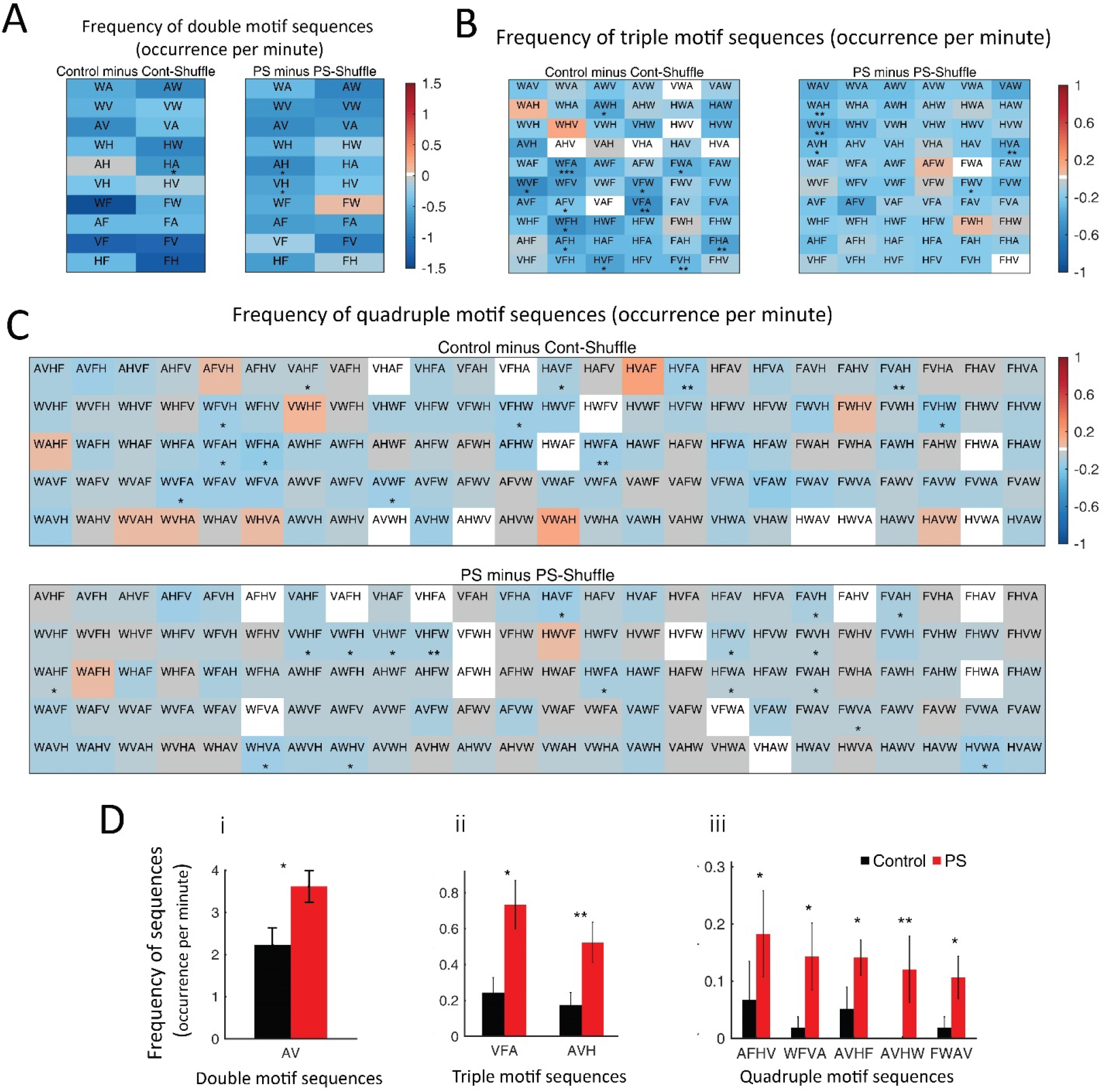
Nonrepetitive motif sequences. **A)** Matrices demonstrate 20 possible nonrepetitive double motif sequences in a 10×2 block. H, F, A, W, and V stand for hindlimb, forelimb, auditory, whisker, and visual sensory motifs, respectively. The letters in each block are a short form of sequences’ names, and the colors reflect changes in the occurrence rate of sequences from a random presentation. In the control group, one cell with asterisks indicates one double sequence (5%) occurred significantly different than random. In the PS group, two cells with asterisks imply two sequences (10%) that occurred differently from shuffle data. **B)** Matrices demonstrate 60 possible nonrepetitive triple motif sequences in a 10×6 block. In the control group, 12 cells with asterisks indicate 12 triple sequences (20%) occurred significantly different than random. In the PS group, 5 sequences (8.3%) occurred differently from shuffle data. **C)** Matrices demonstrate 120 possible nonrepetitive quadruple motif sequences in a 5×24 block. In the control group, 12 quadruple sequences (10%) occurred significantly different than random. In the PS group, 17 sequences (14.1%) occurred differently from shuffle data. **D)** (i) One out of 20 sequences of double combinations of motifs occurred significantly different in control and the PS groups (AV: df=12, sum of ranks=63, p=0.02). (ii) Two out of 60 sequences of triple combinations of motifs occurred significantly different in the two groups (VFA: df=12, sum of ranks=48, p=0.017; AVH: df=12, sum of ranks=64, p=0.008). (iii) Five out of 120 sequences of quadruple combinations of motifs occurred significantly different in the two groups (AFHV: df=12, sum of ranks=61, p=0.032; WFVA: df=12, sum of ranks=61.5, p=0.022; AVHF: df=12, sum of ranks=59.5, p=0.035; AVHW: df=12, sum of ranks=65, p=0.006; FWAV: df=12, sum of ranks=60.5, p=0.027). The remaining sequences not shown in panel **D** did not occur significantly different in the two groups. The number of mice: n=8 control, n=6 PS. Asterisks indicate *p<0.05, **p<0.01, or ***p<0.001.

**Supplementary Table 1.**
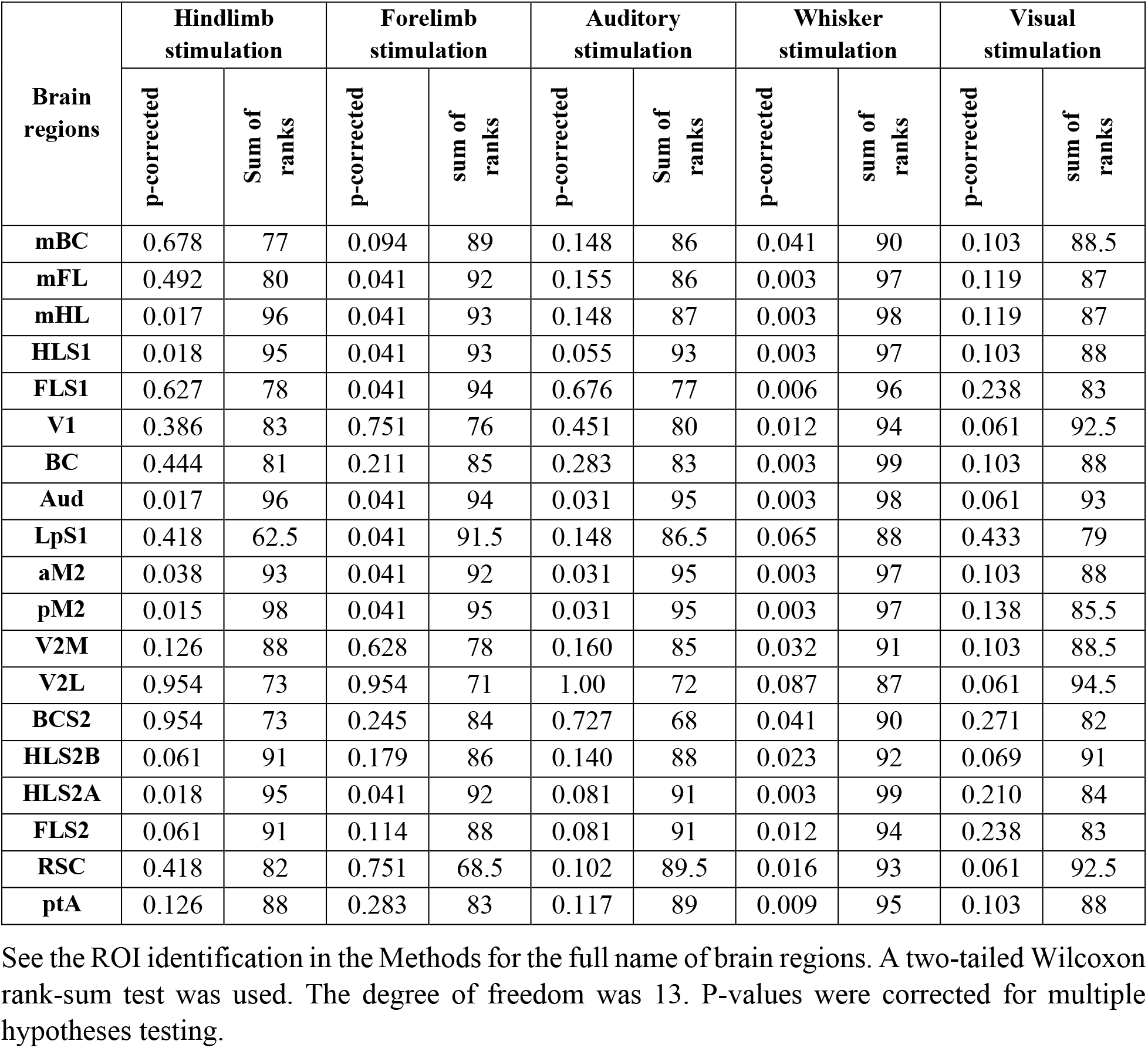
Statistical parameters for the peak amplitude in Supplementary Fig. 1

**Supplementary Table 2.**
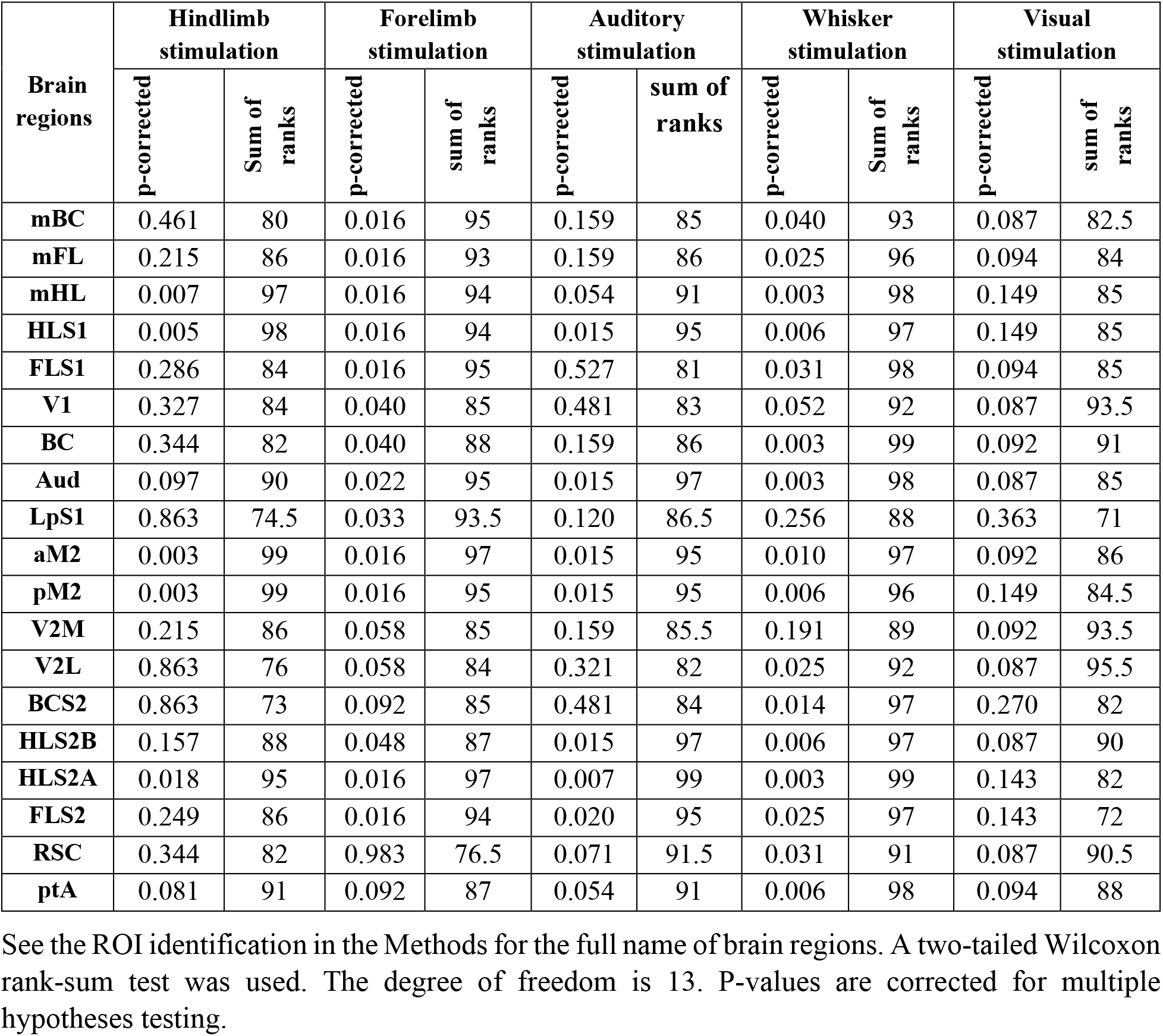
Statistical parameters for the area under the peak in Supplementary Fig. 2

**Supplementary Table 3.** Statistical parameters for the rise time in Supplementary Fig. 3

| Brain regions | Hindlimb stimulation |  | Forelimb stimulation |  | Auditory stimulation |  | Whisker stimulation |  | Visual stimulation |  |
| --- | --- | --- | --- | --- | --- | --- | --- | --- | --- | --- |
|  | p-corrected | Sum of ranks | p-corrected | sum of ranks | p-corrected | sum of ranks | p-corrected | Sum of ranks | p-corrected | sum of ranks |
| mBC | 0.431 | 82 | 0.357 | 85 | 0.649 | 80 | 0.131 | 90.5 | 0.498 | 84.5 |
| mFL | 0.146 | 89 | 0.332 | 88 | 0.735 | 78 | 0.456 | 81.5 | 0.349 | 91 |
| mHL | 0.015 | 97.5 | 0.400 | 83.5 | 0.355 | 86 | 0.086 | 92.5 | 0.498 | 85 |
| HLS1 | 0.043 | 94.5 | 0.201 | 91.5 | 0.355 | 86 | 0.086 | 93.5 | 0.794 | 75 |
| FLS1 | 0.146 | 88 | 0.644 | 78.5 | 0.941 | 73.5 | 0.208 | 87 | 0.498 | 83.5 |
| V1 | 0.431 | 82 | 0.980 | 71.5 | 0.451 | 83 | 0.208 | 87 | 0.794 | 75.5 |
| BC | 0.937 | 74 | 0.357 | 85.5 | 0.866 | 75 | 1.00 | 72 | 0.794 | 69.5 |
| Aud | 0.937 | 73.5 | 0.685 | 77 | 0.866 | 68.5 | 0.326 | 84 | 0.670 | 81 |
| LpS1 | 0.717 | 66 | 0.332 | 87.5 | 0.451 | 83.5 | 0.792 | 75 | 0.498 | 82.5 |
| aM2 | 0.222 | 86 | 0.644 | 80 | 0.201 | 91 | 0.528 | 80 | 0.498 | 84 |
| pM2 | 0.093 | 91.5 | 0.644 | 79 | 0.451 | 83 | 0.086 | 92.5 | 0.794 | 74.5 |
| V2M | 0.954 | 73 | 0.357 | 86.5 | 0.355 | 87.5 | 0.208 | 87 | 0.498 | 88.5 |
| V2L | 0.717 | 78 | 0.644 | 78 | 0.355 | 86 | 0.528 | 79 | 0.794 | 77.5 |
| BCS2 | 0.323 | 84 | 0.704 | 76 | 0.735 | 78.5 | 0.528 | 79 | 0.794 | 76 |
| HLS2B | 0.146 | 88 | 0.400 | 84 | 0.950 | 73 | 0.611 | 77.5 | 0.498 | 83 |
| HLS2A | 0.146 | 88 | 0.644 | 78 | 0.735 | 65 | 0.528 | 79 | 0.794 | 67.5 |
| FLS2 | 0.795 | 76.5 | 0.704 | 68 | 0.866 | 68 | 0.326 | 83.5 | 0.794 | 75 |
| RSC | 0.937 | 70 | 0.644 | 78.5 | 0.201 | 92.5 | 0.286 | 85 | 0.794 | 78 |
| ptA | 0.007 | 99 | 0.083 | 95 | 0.866 | 75 | 0.280 | 89 | 0.794 | 78 |

See the ROI identification in the Methods for the full name of brain regions. A two-tailed Wilcoxon rank-sum test was used. The degree of freedom is 13. P-values are corrected for multiple hypotheses testing.

**Supplementary Table 4.**
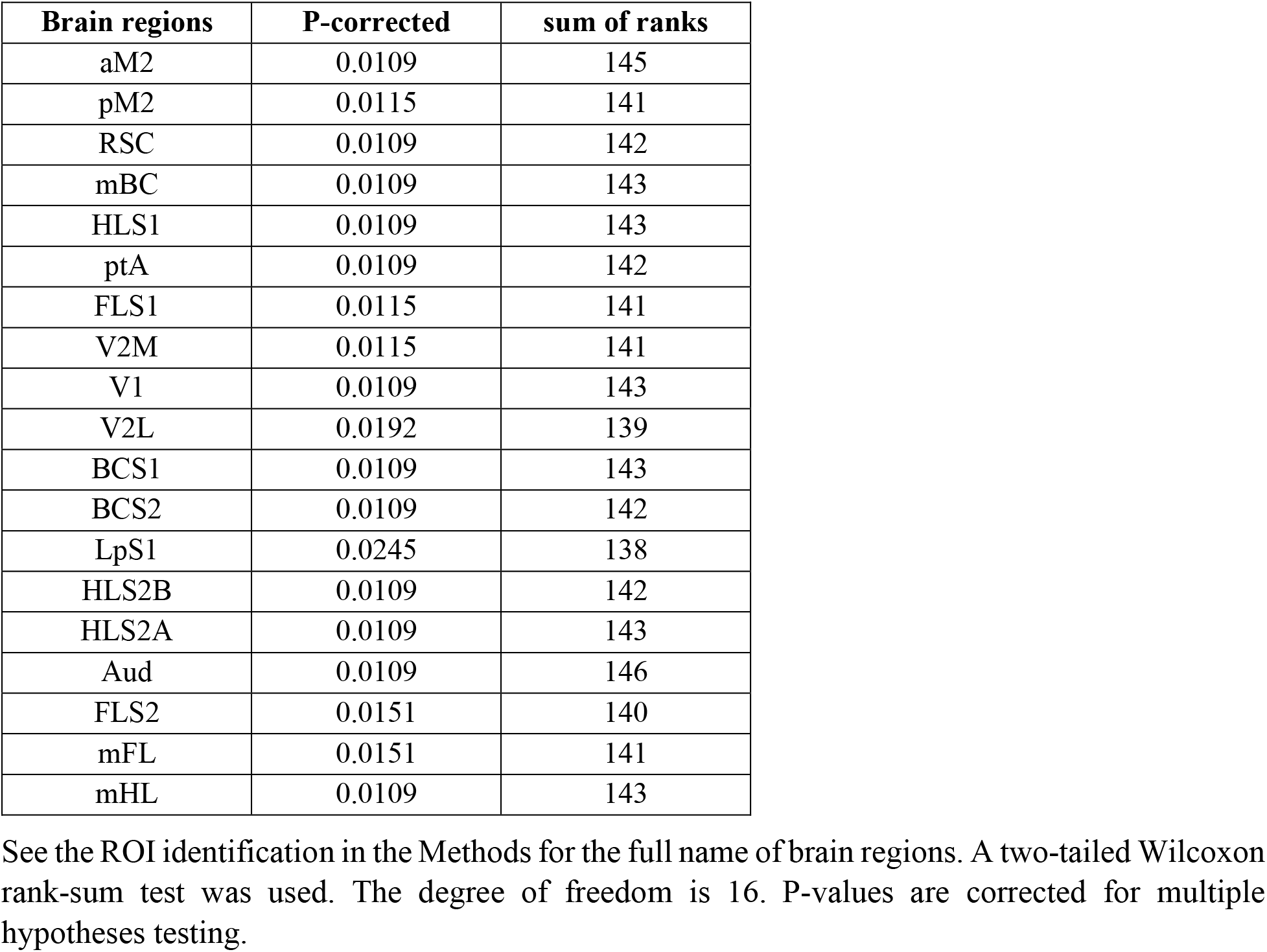
Statistical parameters for the bar graph in Fig. 2B

**Supplementary Table 5.** Statistical parameters from the Wilcoxon rank-sum test for network comparisons

| **Network measures** | **Control-Shuffle vs PS-Shuffle** |  | **Control vs Control-Shuffle** |  | **PS vs PS-Shuffle** |  |
| --- | --- | --- | --- | --- | --- | --- |
|  | p-corrected | Sum of ranks | p-corrected | Sum of ranks | p-corrected | Sum of ranks |
| Efficiency | 0.0063 | 143 | 0.9539 | 152 | 0.5887 | 43 |
| Clustering coefficient | 0.0063 | 143 | 0.9539 | 144 | 0.5887 | 35 |
| Characteristic path | 0.0068 | 86 | 0.9539 | 151.5 | 0.5877 | 34 |
| Betweenness centrality | 0.0063 | 86 | 0.9539 | 156 | 0.2510 | 51 |

Network measures are tested for statistical significance when the correlation matrices are shuffled. Under shuffling, group differences are significant while there is no significant difference in the network measures for a given group, confirming the encoding of network information in weight distributions. A two-tailed Wilcoxon rank-sum test was used. The degree of freedom is 16. P-values are corrected for multiple hypotheses testing.

## References

Abbott, P.W., Gumusoglu, S.B., Bittle, J., Beversdorf, D.Q., Stevens, H.E., 2018. Prenatal stress and genetic risk: How prenatal stress interacts with genetics to alter risk for psychiatric illness. Psychoneuroendocrinology 90, 9–21.

Afrashteh, N., Inayat, S., Bermudez Contreras, E., Luczak, A., McNaughton, B., Mohajerani, M., Spatiotemporal structure of sensory-evoked and spontaneous activity revealed by mesoscale imaging in anesthetized and awake mice. BioRxiv.

Amaya, J.M., Suidgeest, E., Sahut-Barnola, I., Dumontet, T., Montanier, N., Pagès, G., Keller, C., van der Weerd, L., Pereira, A.M., Martinez, A., Meijer, O.C., 2021. Effects of Long-Term Endogenous Corticosteroid Exposure on Brain Volume and Glial Cells in the AdKO Mouse. Frontiers in neuroscience 15, 604103.

Azouz, R., Gray, C.M., 1999. Cellular mechanisms contributing to response variability of cortical neurons in vivo. The Journal of neuroscience : the official journal of the Society for Neuroscience 19, 2209–2223.

Bale, T.L., Baram, T.Z., Brown, A.S., Goldstein, J.M., Insel, T.R., McCarthy, M.M., Nemeroff, C.B., Reyes, T.M., Simerly, R.B., Susser, E.S., Nestler, E.J., 2010. Early life programming and neurodevelopmental disorders. Biol Psychiatry 68, 314–319.

Barttfeld, P., Uhrig, L., Sitt, J.D., Sigman, M., Jarraya, B., Dehaene, S., 2015. Signature of consciousness in the dynamics of resting-state brain activity. Proceedings of the National Academy of Sciences of the United States of America 112, 887–892.

Benjamini, Y., Hochberg, Y., 1995. Controlling the False Discovery Rate: A Practical and Powerful Approach to Multiple Testing. J. R. Statist. Soc. B. 57, 289–300.

Bermudez-Contreras, E., Chekhov, S., Sun, J., Tarnowsky, J., McNaughton, B.L., Mohajerani, M.H., 2018. High-performance, inexpensive setup for simultaneous multisite recording of electrophysiological signals and mesoscale voltage imaging in the mouse cortex. Neurophotonics 5, 025005.

Betzel, R., Griffa, A., Koenigsberger, A., Goñi, J., Thiran, J., Hagmann, P., Sporns, O., 2013. Multi-scale community organization of the human structural connectome and its relationship with resting-state functional connectivity. Network Science 1, 353–373.

Betzel, R.F., Bertolero, M.A., Gordon, E.M., Gratton, C., Dosenbach, N.U.F., Bassett, D.S., 2019. The community structure of functional brain networks exhibits scale-specific patterns of inter- and intra-subject variability. Neuroimage 202, 115990.

Blondel, V., Guillaume, J., Lambiotte, R., Lefebvre, E., 2008. Fast unfolding of communities in large networks. Journal of Statistical Mechanics: Theory and Experiment 2008.

Bock, J., Rether, K., Groger, N., Xie, L., Braun, K., 2014. Perinatal programming of emotional brain circuits: an integrative view from systems to molecules. Front Neurosci 8, 11.

Bock, J., Wainstock, T., Braun, K., Segal, M., 2015. Stress In Utero: Prenatal Programming of Brain Plasticity and Cognition. Biol Psychiatry 78, 315–326.

Bokil, H., Andrews, P., Kulkarni, J.E., Mehta, S., Mitra, P.P., 2010. Chronux: a platform for analyzing neural signals. J Neurosci Methods 192, 146–151.

Bronson, S.L., Bale, T.L., 2014. Prenatal stress-induced increases in placental inflammation and offspring hyperactivity are male-specific and ameliorated by maternal antiinflammatory treatment. Endocrinology 155, 2635–2646.

Buchanan, C.R., Bastin, M.E., Ritchie, S.J., Liewald, D.C., Madole, J.W., Tucker-Drob, E.M., Deary, I.J., Cox, S.R., 2020. The effect of network thresholding and weighting on structural brain networks in the UK Biobank. Neuroimage 211, 116443.

Bullmore, E., Sporns, O., 2009. Complex brain networks: graph theoretical analysis of structural and functional systems. Nat Rev Neurosci 10, 186–198.

Burow, A., Day, H.E., Campeau, S., 2005. A detailed characterization of loud noise stress: Intensity analysis of hypothalamo-pituitary-adrenocortical axis and brain activation. Brain research 1062, 63–73.

Chang, T.Y., Beelen, R., Li, S.F., Chen, T.I., Lin, Y.J., Bao, B.Y., Liu, C.S., 2014. Road traffic noise frequency and prevalent hypertension in Taichung, Taiwan: a cross-sectional study. Environ Health 13, 37.

Charil, A., Laplante, D.P., Vaillancourt, C., King, S., 2010. Prenatal stress and brain development. Brain Res Rev 65, 56–79.

Chen, G.D., Sheppard, A., Salvi, R., 2016. Noise trauma induced plastic changes in brain regions outside the classical auditory pathway. Neuroscience 315, 228–245.

Clancy, B., Finlay, B.L., Darlington, R.B., Anand, K.J., 2007. Extrapolating brain development from experimental species to humans. Neurotoxicology 28, 931–937.

Crossley, N.A., Mechelli, A., Scott, J., Carletti, F., Fox, P.T., McGuire, P., Bullmore, E.T., 2014. The hubs of the human connectome are generally implicated in the anatomy of brain disorders. Brain 137, 2382–2395.

Crudo, A., Petropoulos, S., Moisiadis, V.G., Iqbal, M., Kostaki, A., Machnes, Z., Szyf, M., Matthews, S.G., 2012. Prenatal synthetic glucocorticoid treatment changes DNA methylation states in male organ systems: multigenerational effects. Endocrinology 153, 3269–3283.

Czeh, B., Muller-Keuker, J.I., Rygula, R., Abumaria, N., Hiemke, C., Domenici, E., Fuchs, E., 2007. Chronic social stress inhibits cell proliferation in the adult medial prefrontal cortex: hemispheric asymmetry and reversal by fluoxetine treatment. Neuropsychopharmacology 32, 1490–1503.

de Kloet, E.R., Joels, M., Holsboer, F., 2005. Stress and the brain: from adaptation to disease. Nature reviews. Neuroscience 6, 463–475.

Devilbiss, D.M., Waterhouse, B.D., Berridge, C.W., Valentino, R., 2012. Corticotropin-releasing factor acting at the locus coeruleus disrupts thalamic and cortical sensory-evoked responses. Neuropsychopharmacology : official publication of the American College of Neuropsychopharmacology 37, 2020–2030.

Eggermont, J.J., 2014. Noise and the Brain. Academic Press, Elsevier.

Epskamp, S., Fried, E.I., 2018. A tutorial on regularized partial correlation networks. Psychological methods 23, 617–634.

Estes, M.L., McAllister, A.K., 2016. Maternal immune activation: Implications for neuropsychiatric disorders. Science 353, 772–777.

Favaro, A., Tenconi, E., Degortes, D., Manara, R., Santonastaso, P., 2015. Neural correlates of prenatal stress in young women. Psychol Med 45, 2533–2543.

Ferezou, I., Deneux, T., 2017. Review: How do spontaneous and sensory-evoked activities interact? Neurophotonics 4, 031221.

Fornito, F., Zalesky, A., Bullmore, E., 2016. Fundamentals of Brain Network Analysis. Elsevier: Academic Press.

Friston, K.J., 2011. Functional and effective connectivity: a review. Brain Connect 1, 13–36.

Graham, A.M., Buss, C., Rasmussen, J.M., Rudolph, M.D., Demeter, D.V., Gilmore, J.H., Styner, M., Entringer, S., Wadhwa, P.D., Fair, D.A., 2016. Implications of newborn amygdala connectivity for fear and cognitive development at 6-months-of-age. Dev Cogn Neurosci 18, 12–25.

Graham, A.M., Rasmussen, J.M., Entringer, S., Ben Ward, E., Rudolph, M.D., Gilmore, J.H., Styner, M., Wadhwa, P.D., Fair, D.A., Buss, C., 2019. Maternal Cortisol Concentrations During Pregnancy and Sex-Specific Associations With Neonatal Amygdala Connectivity and Emerging Internalizing Behaviors. Biol Psychiatry 85, 172–181.

Grandjean, J., Canella, C., Anckaerts, C., Ayrancı, G., Bougacha, S., Bienert, T., Buehlmann, D., Coletta, L., Gallino, D., Gass, N., Garin, C.M., Nadkarni, N.A., Hübner, N.S., Karatas, M., Komaki, Y., Kreitz, S., Mandino, F., Mechling, A.E., Sato, C., Sauer, K., Shah, D., Strobelt, S., Takata, N., Wank, I., Wu, T., Yahata, N., Yeow, L.Y., Yee, Y., Aoki, I., Chakravarty, M.M., Chang, W.T., Dhenain, M., von Elverfeldt, D., Harsan, L.A., Hess, A., Jiang, T., Keliris, G.A., Lerch, J.P., Meyer-Lindenberg, A., Okano, H., Rudin, M., Sartorius, A., Van der Linden, A., Verhoye, M., Weber-Fahr, W., Wenderoth, N., Zerbi, V., Gozzi, A., 2020. Common functional networks in the mouse brain revealed by multi-centre resting-state fMRI analysis. Neuroimage 205, 116278.

Grandjean, J., Zerbi, V., Balsters, J.H., Wenderoth, N., Rudin, M., 2017. Structural Basis of Large-Scale Functional Connectivity in the Mouse. The Journal of neuroscience : the official journal of the Society for Neuroscience 37, 8092–8101.

Han, F., Caporale, N., Dan, Y., 2008. Reverberation of recent visual experience in spontaneous cortical waves. Neuron 60, 321–327.

Han, K., Min, J., Lee, M., Kang, B.M., Park, T., Hahn, J., Yei, J., Lee, J., Woo, J., Lee, C.J., Kim, S.G., Suh, M., 2019. Neurovascular Coupling under Chronic Stress Is Modified by Altered GABAergic Interneuron Activity. The Journal of neuroscience : the official journal of the Society for Neuroscience 39, 10081–10095.

Handa, R.J., Weiser, M.J., 2014. Gonadal steroid hormones and the hypothalamo-pituitary-adrenal axis. Front Neuroendocrinol 35, 197–220.

Heffner, H.E., Heffner, R.S., 2007. Hearing ranges of laboratory animals. J Am Assoc Lab Anim Sci 46, 20–22.

Hilgetag, C.C., Burns, G.A., O’Neill, M.A., Scannell, J.W., Young, M.P., 2000. Anatomical connectivity defines the organization of clusters of cortical areas in the macaque monkey and the cat. Philos Trans R Soc Lond B Biol Sci 355, 91–110.

Huizink, A.C., Mulder, E.J., Buitelaar, J.K., 2004. Prenatal stress and risk for psychopathology: specific effects or induction of general susceptibility? Psychol Bull 130, 115–142.

Jafari, Z., Faraji, J., Mirza Agha, B., Metz, G.A.S., Kolb, B.E., Mohajerani, M.H., 2017a. The Adverse Effects of Auditory Stress on Mouse Uterus Receptivity and Behaviour. Sci Rep 7, 4720.

Jafari, Z., Kolb, B.E., Mohajerani, M.H., 2018a. Chronic traffic noise stress accelerates brain impairment and cognitive decline in mice. Exp Neurol 308, 1–12.

Jafari, Z., Kolb, B.E., Mohajerani, M.H., 2020a. Life-Course Contribution of Prenatal Stress in Regulating the Neural Modulation Network Underlying the Prepulse Inhibition of the Acoustic Startle Reflex in Male Alzheimer’s Disease Mice. Cerebral cortex (New York, N.Y. : 1991) 30, 311–325.

Jafari, Z., Kolb, B.E., Mohajerani, M.H., 2020b. Noise exposure accelerates the risk of cognitive impairment and Alzheimer’s disease: Adulthood, gestational, and prenatal mechanistic evidence from animal studies. Neuroscience and biobehavioral reviews 117, 110–128.

Jafari, Z., Mehla, J., Afrashteh, N., Kolb, B.E., Mohajerani, M.H., 2017b. Corticosterone response to gestational stress and postpartum memory function in mice. PloS one 12, e0180306.

Jafari, Z., Mehla, J., Kolb, B.E., Mohajerani, M.H., 2018b. Gestational Stress Augments Postpartum β-Amyloid Pathology and Cognitive Decline in a Mouse Model of Alzheimer’s Disease. Cerebral cortex (New York, N.Y. : 1991), 1–13.

Jafari, Z., Okuma, M., Karem, H., Mehla, J., Kolb, B.E., Mohajerani, M.H., 2019. Prenatal noise stress aggravates cognitive decline and the onset and progression of beta amyloid pathology in a mouse model of Alzheimer’s disease. Neurobiology of aging 77, 66–86.

Jensen Pena, C., Monk, C., Champagne, F.A., 2012. Epigenetic effects of prenatal stress on 11beta-hydroxysteroid dehydrogenase-2 in the placenta and fetal brain. PloS one 7, e39791.

Joels, M., Karst, H., Alfarez, D., Heine, V.M., Qin, Y., van Riel, E., Verkuyl, M., Lucassen, P.J., Krugers, H.J., 2004. Effects of chronic stress on structure and cell function in rat hippocampus and hypothalamus. Stress (Amsterdam, Netherlands) 7, 221–231.

Joels, M., Pu, Z., Wiegert, O., Oitzl, M.S., Krugers, H.J., 2006. Learning under stress: how does it work? Trends in cognitive sciences 10, 152–158.

Kazeminejad, A., Sotero, R.C., 2020. The Importance of Anti-correlations in Graph Theory Based Classification of Autism Spectrum Disorder. Frontiers in neuroscience 14, 676.

Kenet, T., Bibitchkov, D., Tsodyks, M., Grinvald, A., Arieli, A., 2003. Spontaneously emerging cortical representations of visual attributes. Nature 425, 954–956.

Kim, D.J., Davis, E.P., Sandman, C.A., Sporns, O., O’Donnell, B.F., Buss, C., Hetrick, W.P., 2017. Prenatal Maternal Cortisol Has Sex-Specific Associations with Child Brain Network Properties. Cerebral cortex (New York, N.Y. : 1991) 27, 5230–5241.

Kim, E.J., Pellman, B., Kim, J.J., 2015. Stress effects on the hippocampus: a critical review. Learn Mem 22, 411–416.

Kinney, D.K., Munir, K.M., Crowley, D.J., Miller, A.M., 2008. Prenatal stress and risk for autism. Neuroscience and biobehavioral reviews 32, 1519–1532.

Kolb, B., Mychasiuk, R., Muhammad, A., Gibb, R., 2013. Brain plasticity in the developing brain. Prog Brain Res 207, 35–64.

Kolb, B., Mychasiuk, R., Muhammad, A., Li, Y., Frost, D.O., Gibb, R., 2012. Experience and the developing prefrontal cortex. Proceedings of the National Academy of Sciences of the United States of America 109 Suppl 2, 17186–17193.

Kong, Y., Eippert, F., Beckmann, C.F., Andersson, J., Finsterbusch, J., Buchel, C., Tracey, I., Brooks, J.C., 2014. Intrinsically organized resting state networks in the human spinal cord. Proceedings of the National Academy of Sciences of the United States of America 111, 18067–18072.

Kyweriga, M., Sun, J., Wang, S., Kline, R., Mohajerani, M.H., 2017. A Large Lateral Craniotomy Procedure for Mesoscale Wide-field Optical Imaging of Brain Activity. J Vis Exp.

Lancichinetti, A., Fortunato, S., 2012. Consensus clustering in complex networks. Sci Rep 2, 336.

Liska, A., Galbusera, A., Schwarz, A.J., Gozzi, A., 2015. Functional connectivity hubs of the mouse brain. Neuroimage 115, 281–291.

Liu, J., Chen, X., Lewis, G., 2011. Childhood internalizing behaviour: analysis and implications. J Psychiatr Ment Health Nurs 18, 884–894.

Lussier, S.J., Stevens, H.E., 2016. Delays in GABAergic interneuron development and behavioral inhibition after prenatal stress. Dev Neurobiol 76, 1078–1091.

Marecková, K., Klasnja, A., Bencurova, P., Andrýsková, L., Brázdil, M., Paus, T., 2019. Prenatal Stress, Mood, and Gray Matter Volume in Young Adulthood. Cerebral cortex (New York, N.Y. : 1991) 29, 1244–1250.

Martin, E.A., Hlinka, J., Meinke, A., Dechterenko, F., Tintera, J., Oliver, I., Davidsen, J., 2017. Network Inference and Maximum Entropy Estimation on Information Diagrams. Sci Rep 7, 7062.

Maselko, J., Sikander, S., Bhalotra, S., Bangash, O., Ganga, N., Mukherjee, S., Egger, H., Franz, L., Bibi, A., Liaqat, R., Kanwal, M., Abbasi, T., Noor, M., Ameen, N., Rahman, A., 2015. Effect of an early perinatal depression intervention on long-term child development outcomes: follow-up of the Thinking Healthy Programme randomised controlled trial. Lancet Psychiatry 2, 609–617.

Maxwell, C.R., Ehrlichman, R.S., Liang, Y., Gettes, D.R., Evans, D.L., Kanes, S.J., Abel, T., Karp, J., Siegel, S.J., 2006. Corticosterone modulates auditory gating in mouse. Neuropsychopharmacology 31, 897–903.

McCormick, D.A., 1999. Spontaneous activity: signal or noise? Science 285, 541–543.

McEwen, B.S., 2004. Protection and damage from acute and chronic stress: allostasis and allostatic overload and relevance to the pathophysiology of psychiatric disorders. Annals of the New York Academy of Sciences 1032, 1–7.

McGirr, A., LeDue, J., Chan, A.W., Boyd, J.D., Metzak, P.D., Murphy, T.H., 2020. Stress impacts sensory variability through cortical sensory activity motifs. Transl Psychiatry 10, 20.

Meyer, U., Murray, P.J., Urwyler, A., Yee, B.K., Schedlowski, M., Feldon, J., 2008. Adult behavioral and pharmacological dysfunctions following disruption of the fetal brain balance between pro-inflammatory and IL-10-mediated anti-inflammatory signaling. Mol Psychiatry 13, 208–221.

Mohajerani, M.H., Chan, A.W., Mohsenvand, M., LeDue, J., Liu, R., McVea, D.A., Boyd, J.D., Wang, Y.T., Reimers, M., Murphy, T.H., 2013. Spontaneous cortical activity alternates between motifs defined by regional axonal projections. Nature neuroscience 16, 1426–1435.

Mohajerani, M.H., McVea, D.A., Fingas, M., Murphy, T.H., 2010. Mirrored bilateral slow-wave cortical activity within local circuits revealed by fast bihemispheric voltage-sensitive dye imaging in anesthetized and awake mice. The Journal of neuroscience : the official journal of the Society for Neuroscience 30, 3745–3751.

Mychasiuk, R., Gibb, R., Kolb, B., 2012. Prenatal stress alters dendritic morphology and synaptic connectivity in the prefrontal cortex and hippocampus of developing offspring. Synapse 66, 308–314.

Nasiriavanaki, M., Xia, J., Wan, H., Bauer, A.Q., Culver, J.P., Wang, L.V., 2014. High-resolution photoacoustic tomography of resting-state functional connectivity in the mouse brain. Proceedings of the National Academy of Sciences of the United States of America 111, 21–26.

Negron-Oyarzo, I., Neira, D., Espinosa, N., Fuentealba, P., Aboitiz, F., 2015. Prenatal Stress Produces Persistence of Remote Memory and Disrupts Functional Connectivity in the Hippocampal-Prefrontal Cortex Axis. Cerebral cortex (New York, N.Y. : 1991) 25, 3132–3143.

Newman, M.E., 2006. Modularity and community structure in networks. Proceedings of the National Academy of Sciences of the United States of America 103, 8577–8582.

O’Donnell, K.J., Bugge Jensen, A., Freeman, L., Khalife, N., O’Connor, T.G., Glover, V., 2012. Maternal prenatal anxiety and downregulation of placental 11beta-HSD2. Psychoneuroendocrinology 37, 818–826.

Oliver, I., Hlinka, J., Kopal, J., Davidsen, J., 2019. Quantifying the Variability in Resting-State Networks. Entropy 21, 882.

Pavlovska-Teglia, G., Stodulski, G., Svendsen, L., Dalton, K., Hau, J., 1995. Effect of oral corticosterone administration on locomotor development of neonatal and juvenile rats. Exp Physiol 80, 469–475.

Paxinos, G., and Franklin, KBJ., 2001. The Mouse Brain in Stereotaxic Coordinates. CA: Academic Press, San Diego.

Power, J.D., Schlaggar, B.L., Lessov-Schlaggar, C.N., Petersen, S.E., 2013. Evidence for hubs in human functional brain networks. Neuron 79, 798–813.

Raichle, M.E., 2010. Two views of brain function. Trends Cogn Sci 14, 180–190.

Reynolds, R.M., 2013. Glucocorticoid excess and the developmental origins of disease: two decades of testing the hypothesis--2012 Curt Richter Award Winner. Psychoneuroendocrinology 38, 1–11.

Ringach, D.L., 2009. Spontaneous and driven cortical activity: implications for computation. Curr Opin Neurobiol 19, 439–444.

Rubin, L.P., 2016. Maternal and pediatric health and disease: integrating biopsychosocial models and epigenetics. Pediatr Res 79, 127–135.

Rubinov, M., Sporns, O., 2010. Complex network measures of brain connectivity: uses and interpretations. Neuroimage 52, 1059–1069.

Salas, M., Schapiro, S., 1970. Hormonal influences upon the maturation of the rat brain’s responsiveness to sensory stimuli. Physiol Behav 5, 7–11.

Sapolsky, R.M., Romero, L.M., Munck, A.U., 2000. How do glucocorticoids influence stress responses? Integrating permissive, suppressive, stimulatory, and preparative actions. Endocr Rev 21, 55–89.

Scheinost, D., Kwon, S.H., Lacadie, C., Sze, G., Sinha, R., Constable, R.T., Ment, L.R., 2016. Prenatal stress alters amygdala functional connectivity in preterm neonates. Neuroimage Clin 12, 381–388.

Scheinost, D., Sinha, R., Cross, S.N., Kwon, S.H., Sze, G., Constable, R.T., Ment, L.R., 2017. Does prenatal stress alter the developing connectome? Pediatr Res 81, 214–226.

Scheinost, D., Spann, M.N., McDonough, L., Peterson, B.S., Monk, C., 2020. Associations between different dimensions of prenatal distress, neonatal hippocampal connectivity, and infant memory. Neuropsychopharmacology : official publication of the American College of Neuropsychopharmacology 45, 1272–1279.

Schwarz, A.J., McGonigle, J., 2011. Negative edges and soft thresholding in complex network analysis of resting state functional connectivity data. NeuroImage 55, 1132–1146.

Shoham, D., Glaser, D.E., Arieli, A., Kenet, T., Wijnbergen, C., Toledo, Y., Hildesheim, R., Grinvald, A., 1999. Imaging cortical dynamics at high spatial and temporal resolution with novel blue voltage-sensitive dyes. Neuron 24, 791–802.

Smith, G.B., Hein, B., Whitney, D.E., Fitzpatrick, D., Kaschube, M., 2018. Distributed network interactions and their emergence in developing neocortex. Nature neuroscience 21, 1600–1608.

Smith, S.M., Miller, K.L., Salimi-Khorshidi, G., Webster, M., Beckmann, C.F., Nichols, T.E., Ramsey, J.D., Woolrich, M.W., 2011. Network modelling methods for FMRI. NeuroImage 54, 875–891.

Smith, S.M., Vidaurre, D., Beckmann, C.F., Glasser, M.F., Jenkinson, M., Miller, K.L., Nichols, T.E., Robinson, E.C., Salimi-Khorshidi, G., Woolrich, M.W., Barch, D.M., Uğurbil, K., Van Essen, D.C., 2013. Functional connectomics from resting-state fMRI. Trends in cognitive sciences 17, 666–682.

Spann, M.N., Monk, C., Scheinost, D., Peterson, B.S., 2018. Maternal Immune Activation During the Third Trimester Is Associated with Neonatal Functional Connectivity of the Salience Network and Fetal to Toddler Behavior. The Journal of neuroscience : the official journal of the Society for Neuroscience 38, 2877–2886.

Sporns, O., 2011. The non-random brain: efficiency, economy, and complex dynamics. Frontiers in computational neuroscience 5, 5.

Sporns, O., 2014. Towards network substrates of brain disorders. Brain 137, 2117–2118.

Sporns, O., Betzel, R.F., 2016. Modular Brain Networks. Annual review of psychology 67, 613–640.

Sporns, O., Honey, C.J., Kotter, R., 2007. Identification and classification of hubs in brain networks. PloS one 2, e1049.

Sporns, O., Zwi, J.D., 2004. The small world of the cerebral cortex. Neuroinformatics 2, 145–162.

Stevens, K.E., Bullock, A.E., Collins, A.C., 2001. Chronic corticosterone treatment alters sensory gating in C3H mice. Pharmacology, biochemistry, and behavior 69, 359–366.

Thomason, M.E., Brown, J.A., Dassanayake, M.T., Shastri, R., Marusak, H.A., Hernandez-Andrade, E., Yeo, L., Mody, S., Berman, S., Hassan, S.S., Romero, R., 2014. Intrinsic functional brain architecture derived from graph theoretical analysis in the human fetus. PloS one 9, e94423.

Turesky, T.K., Jensen, S.K.G., Yu, X., Kumar, S., Wang, Y., Sliva, D.D., Gagoski, B., Sanfilippo, J., Zollei, L., Boyd, E., Haque, R., Hafiz Kakon, S., Islam, N., Petri, W.A., Jr., Nelson, C.A., Gaab, N., 2019. The relationship between biological and psychosocial risk factors and resting-state functional connectivity in 2-month-old Bangladeshi infants: A feasibility and pilot study. Dev Sci 22, e12841.

Turkheimer, F.E., Leech, R., Expert, P., Lord, L.D., Vernon, A.C., 2015. The brain’s code and its canonical computational motifs. From sensory cortex to the default mode network: A multi-scale model of brain function in health and disease. Neuroscience and biobehavioral reviews 55, 211–222.

van den Bergh, B.R.H., Dahnke, R., Mennes, M., 2018. Prenatal stress and the developing brain: Risks for neurodevelopmental disorders. Dev Psychopathol 30, 743–762.

Van den Bergh, B.R.H., van den Heuvel, M.I., Lahti, M., Braeken, M., de Rooij, S.R., Entringer, S., Hoyer, D., Roseboom, T., Raikkonen, K., King, S., Schwab, M., 2017. Prenatal developmental origins of behavior and mental health: The influence of maternal stress in pregnancy. Neuroscience and biobehavioral reviews.

van Wijk, B.C., Stam, C.J., Daffertshofer, A., 2010. Comparing brain networks of different size and connectivity density using graph theory. PloS one 5, e13701.

Weinstock, M., 2017. Prenatal stressors in rodents: Effects on behavior. Neurobiol Stress 6, 3–13.

White, D.R., Boettcher, F.A., Miles, L.R., Gratton, M.A., 1998. Effectiveness of intermittent and continuous acoustic stimulation in preventing noise-induced hearing and hair cell loss. J Acoust Soc Am 103, 1566–1572.

Wilson, M.A., Grillo, C.A., Fadel, J.R., Reagan, L.P., 2015. Stress as a one-armed bandit: Differential effects of stress paradigms on the morphology, neurochemistry and behavior in the rodent amygdala. Neurobiol Stress 1, 195–208.

Wu, X.Y., Hu, Y.T., Guo, L., Lu, J., Zhu, Q.B., Yu, E., Wu, J.L., Shi, L.G., Huang, M.L., Bao, A.M., 2015. Effect of pentobarbital and isoflurane on acute stress response in rat. Physiology & behavior 145, 118–121.

Xiao, D., Vanni, M.P., Mitelut, C.C., Chan, A.W., LeDue, J.M., Xie, Y., Chen, A.C., Swindale, N.V., Murphy, T.H., 2017. Mapping cortical mesoscopic networks of single spiking cortical or sub-cortical neurons. eLife 6.

Zaretsky, M.V., Alexander, J.M., Byrd, W., Bawdon, R.E., 2004. Transfer of inflammatory cytokines across the placenta. Obstet Gynecol 103, 546–550.

